# Light responses of melanopsin-expressing ganglion cells in the foetal mammalian retina

**DOI:** 10.1101/675702

**Authors:** Jan Verweij, Shawnta Y. Chaney, Derek Bredl, Shruti Vemaraju, Gabriele M. König, Evi Kostenis, Richard A. Lang, David R. Copenhagen

**Affiliations:** Departments of Ophthalmology and Physiology, University of California, San Francisco, San Francisco, California Box 0444, 94158, USA; Department of Ophthalmology, University of California, San Francisco, San Francisco, California 94143, USA; The Visual Systems Group, Abrahamson Pediatric Eye Institute, Division of Pediatric Ophthalmology, Cincinnati Children’s Hospital Medical Center, Cincinnati, OH 45229, USA; Division of Developmental Biology, University of Cincinnati College of Medicine, Cincinnati, OH 45229, USA; Department of Ophthalmology, University of Cincinnati College of Medicine, Cincinnati, OH 45229, USA; Molecular, Cellular and Pharmacobiology Section, Institute of Pharmaceutical Biology, University of Bonn, 53115 Bonn, Germany

## Abstract

Sensory stimulation plays a critical role in the maturation of sensory organs and systems. For example, when deprived of light before birth, foetal mouse pups *in utero* exhibit altered ocular vascular development. Normal vascular development depends on light excitation of melanopsin, a non-rod, non-cone photopigment that is expressed in a subset of ganglion cells (mRGCs) in the retina. However, there is no direct evidence that mRGCs in foetal eyes are light-responsive. Very little is known about how light absorption leads to excitation in these foetal neurons. Using mRGC-specific expression of the calcium indicators GCaMP3 and GCaMP6, we report that foetal mouse mRGCs respond to light as early as 4 days before birth. Further, two distinct G_q/11_-G protein family antagonists, FR9000359 and YM-254890, abolish these light responses. TTX, a blocker of voltage-activated sodium channels, reversibly represses light responses, and FPL6417 and L-cis-diltiazem, which modify L-type calcium channels, respectively increase and reduce light responses. Electrophysiological patch pipette recordings show that embryonic mRGCs respond to light of intensity as low as 2.9 × 10^12^ photons/cm^2^/s. The present findings demonstrate a heretofore unproven but postulated light sensitivity in the retinas of foetal mice and identify the transduction pathways involved. Surprisingly, mRGCs do not function as completely independent photoreceptors but are electrotonically coupled with other mRGCs. Given that melanopsin is expressed in foetal human retinas, these findings support the idea that the eyes of foetal and early preterm infants are likely to exhibit functional photosensitivity.

**Key points:**

- Melanopsin is a light-excitable photopigment expressed in a subset of ganglion cell neurons (mRGCs) in the retinas of many different species of vertebrates. In mature animals, light activation of mRGCs modulates many visual adaptive functions including pupil constriction, entrainment of circadian rhythms, mood and learning. In neonatal pups at ages prior to the developmental onset of visual signalling from rods and cones, melanopsin cells mediate photoaversive behaviour. In foetal pups, light activation of melanopsin cells accelerates maturation of the ocular vasculature. Here, we describe and physiologically characterize the light responses of melanopsin ganglion cells in the retinas of foetal pups.
- MRGCs in embryonic retinas respond to light at least four days prior to birth and exhibit responses to light of intensity as low as 3 × 10^12^ photons/cm^2^/s.
- Phototransduction mechanisms include melanopsin activation of G_q/11_ – G proteins, voltage-activated sodium currents, and voltage-gated L-type calcium currents.
- MRGCs are electrotonically coupled to other mRGCs in foetal retinas.
- We propose that melanopsin-expressing ganglion cells are excited by light while *in utero* and that this excitation relies, for the most part, on phototransduction pathways that have been described in postnatal retinas. Furthermore, we propose that foetal mRGCs have the requisite properties to modulate light-regulated maturation of the ocular vasculature and, perhaps, the development of visual pathways.

## Introduction

Genetic programmes instruct the initial development of all organisms. Innate genetic programming also regulates the maturation of the eye and the visual system. However, light stimulation of maturing visual circuits also plays an important role in refining patterns of neural connectivity and establishing various types of functionality (Breedlove, 2017). Most previous studies have concentrated on the visually mediated refinement of visual centres at ages after rods and cones in the retina begin responding to light to generate visual signals. However, a unique class of intrinsically photosensitive melanopsin-expressing retinal ganglion cells (mRGCs) that can generate visual signals in the absence of input from rods and cones has been found in adult mammalian eyes (Berson et al., 2002; Do and Yau, 2010). In the eyes of foetal mice, markers of melanopsin expression can be observed 9 days before birth and 19 days before the emergence of visual signalling from rods and cones. In human eyes, markers of melanopsin expression appear at 8.5 weeks post-conception (31 weeks before birth) (Tarttelin et al., 2003). The presence of genetic and immunohistological markers for melanopsin raises the possibility that these mRGCs are intrinsically photosensitive and that light-mediated signalling may occur in embryonic eyes. Thus, ambient light from the pregnant mother’s environment that penetrates to the eyes of foetuses might have the ability to control some aspects of foetal development. Indeed, mouse foetuses that are light-deprived during the final four days of gestation exhibit abnormal postnatal ocular vascular development (Rao et al., 2013). Moreover, genetic deletion of melanopsin in foetal pups mimics the effects of light deprivation (Rao et al., 2013). This finding suggests that ambient light passing through the abdominal wall of the pregnant dam can activate melanopsin-expressing cells in foetal eyes and trigger normal development of the ocular vasculature. A strong supporting argument in favour of this hypothesis would be physiological evidence that mRGCs in the embryonic retina are light-responsive. We report here that foetal mRGCs do respond to light.

Rods and cones in vertebrate retinas utilize opsin-based photopigments to capture light. Rod and cone opsins couple to G-proteins belonging to the G_t_ (transducin) family to generate photoresponses (Arshavsky et al., 2002). The photopigments in invertebrate photoreceptors, unlike those in vertebrates, couple to non-transducin G-proteins. Rhodopsin in *Drosophila* photoreceptors couples to G-proteins of the Gq family (Katz and Minke, 2009). Transduction of light stimuli by melanopsin in postnatal mammalian retinas appears not to be coupled via G_t_, but the identity of the G-protein that mediates phototransduction remains controversial. Several studies have demonstrated that mRGCs in postnatal mammals express one or more of four different alpha subunits, *Gnaq*, *Gna11*, *Gna1* and *Gna15*, that are members of the *G_q/11_* gene family (Chew et al., 2014; Graham et al., 2008; Hughes et al., 2015). Some studies have suggested that melanopsin phototransduction is predominantly mediated by *Gnaq/11* coupling (Hughes et al., 2015; Panda et al., 2005; Qiu et al., 2005). However, other reports concluded that melanopsin functions independently of *Gnaq* and Gna*11* (Chew et al., 2014). In addition, in expression systems, melanopsin can be coupled to G_i/o_ (Bailes and Lucas, 2013) and to *G*_t_ (Newman et al. 2003). Although the identification of melanopsin-G-protein coupling in postnatal mice is somewhat controversial, a very recent study demonstrated that melanopsin-*Gnaq* coupling is necessary and sufficient to trigger normal vascular development in the eyes of foetal mice (Vemaraju et al. 2019). Direct measurement of light activity in mRGCs and pharmacological intervention testing whether G_q/11_ mediates the light responses of individual mRGCs in foetal retinas and not merely ocular vascular development remains to be done.

We report here, using genetically encoded calcium indicator proteins expressed in melanopsin cells, that embryonic mRGCs are light-responsive. We further show that administration of the G_q/11_ antagonists FR900359 and YM-254890 strongly attenuates these light responses, supporting a role for melanopsin-mediated G _q/11_ -coupled signalling in phototransduction. These findings convincingly establish that the melanopsin-expressing ganglion cells in retinas of foetal mice as young as embryonic day 15.5 are light-responsive. Given that the *Opn4* gene is expressed in human eyes at gestational week 9 (Tartellin et al. 2003), these results also raise the very interesting possibility that light sensitivity may be conserved in the human foetus and that it may endow the foetal eye with previously unrecognized photosensitivity.

## Methods

### Animals

The experiments were performed in accordance with the policy of The Journal of Physiology and the rules and regulations set forth in the NIH Guidelines for the Care and Use of Laboratory Animals. The Institutional Animal Care and Use Committee of the University of California, San Francisco (Animal Welfare Assurance Number: A3400-01) approved the experiments. The experiments also conformed to the Society for Neuroscience Policy on the Use of Animals in Neuroscience Research. The mice were housed in an AALAC-accredited pathogen-free animal facility at UCSF with ad libitum access to food and water. The experiments were performed on mice of either sex with an age ranging from embryonic day 15.5 (E15.5) to postnatal day 2 (P2). The mice were housed on a 12-h light-dark cycle with lights on at 7 AM and off at 7 PM. The following mouse strains were used to breed the experimental animals in this study: *Opn4^cre^* knock-in mice with Cre recombinase under the control of the endogenous melanopsin promoter (Ecker et al., 2010); *Opn4^GFP^* transgenic mice with the green fluorescent protein gene driven by the melanopsin promoter (Schmidt et al., 2008); Ai14 strain mice with the floxed tdTomato gene in the rosa26 locus (Madisen et al., 2010); C57Bl/6J wild-type mice (Jackson Laboratory, Sacramento, CA); melanopsin knockout mice (Panda et al., 2003); and finally Ai38(B6.Cg-*Gt(ROSA)26Sor^tm38(CAG-GCaMP3)Hze^*/J) and AI96 (B6;129S6-Gt(ROSA)26Sor^tm96(CAG-GCaMP6s)Hze^/J) strains with the floxed GCaMP3 and GCaMP6 genes, respectively, which encode two different calcium sensors, in the rosa26 locus (Jackson Laboratory, Sacramento, CA; Zariwala et al., 2012). Genotyping of *Opn4^Cre^* (Ecker et al., 2010), *Opn4^−^* (Panda et al., 2002), *Ai14* (Madisen et al., 2010) and *Ai38* (Zariwala et al., 2012) mice was performed as described in the referenced articles.

### Tissue preparation

Pregnant females used in the calcium measurements were sacrificed by CO_2_ inhalation followed by cervical dislocation. For the electrophysiological experiments, pregnant females were sacrificed by cervical dislocation. Following laparotomy, the uterine horns were removed and transferred to cold ACSF (119 mM NaCl, 24 mM NaHCO_3_, 1.25 mM NaH_2_PO_4_, 2.5 mM KCl, 2.5 mM CaCl_2_, 1.5 mM MgSO_4_, 10 mM glucose, 2 mM Na pyruvate, and 2 mM Na lactate) that had been previously bubbled with carbogen (95%O_2_/5% CO_2_). The foetuses were dissected free of their extra-embryonic membranes and decapitated, and the eyes were removed and placed in carbogen-bubbled ACSF. For calcium imaging, the retina was freed from each eye under a dissecting microscope, cut radially and teased, photoreceptor side down, onto a small piece of plastic film; it was then flattened with a brush consisting of a single cat whisker while the plastic film was dragged out of the solution on a microscope slide “boat ramp”. The plastic film was lowered upside down onto a polylysine-coated (0.1 mg/mL) glass coverslip (Number 1, round, 12-mm diameter), and the plastic film was floated off the coverslip using a drop of ACSF. These steps were performed under ambient room light at room temperature (RT). The coverslips were kept in carbogen-bubbled ACSF (RT) in darkness until they were mounted in the recording chamber. For each experiment, a coverslip was placed in the recording chamber (Warner Instruments, Hamden, CT) and perfused with carbogen-bubbled ACSF at ~35°C flowing at 0.4 to 0.8 mL per minute. For patch-clamping, retinas were dissected and kept in carbogen-bubbled ACSF. Immediately before each experiment, a radially cut retina was teased ganglion cell side down onto a plastic film using a single cat whisker brush, dragged up the “boat” ramp, and lowered photoreceptor side down onto a coated coverslip. After floating off the plastic film, the coverslip was mounted in the preparation chamber.

### Patch clamping

The perfused recording chamber was mounted on the stage of an upright Zeiss Axioskop FS microscope equipped with infrared DIC optics and epifluorescence (470/40 excitation filter and 525/50 emission filter). An additional optical channel transmitted excitation from a high-wattage 480 nm LED (Sutter Instruments, Novato, CA). A factory-calibrated radiometer (Model S370, UDT Instruments, San Diego, CA) was used to measure the incident irradiance at the retina. Recordings were performed in ACSF heated to ~34°C using 3 stages of heating (a lab-designed warming blanket near the supply reservoirs followed by a feedback-controlled inline heater and a coverslip preparation chamber with a heated bottom (Cell MicroControls, Norfolk, VA) with flow at approximately 1 mL/s. Patch clamp recordings were performed using an EPC-10 amplifier controlled by “PatchMaster” software (Heka Instruments, Holliston, MA). Glass pipettes were formed from Sutter glass (BF150-110) on a Sutter P-97 electrode puller (Sutter Instruments, Novato, CA). The pipettes were filled with 1 of 3 different patch pipette solutions. The standard solution contained (in mM) 120 K gluconate, 5 NaCl, 4 KCl, 2 EGTA, 10 HEPES, 0.3 NaGTP, 4 MgATP, 7 phosphocreatine, and Tris-KOH (pH 7.3) and had an impedance of 4-6 MOhm. The Alexa pipette solution contained (in mM) 116 K gluconate, 2 EGTA, 10 HEPES, 0.3 NaGTP, 4 MgATP, 7 phosphocreatine tris, neurobiotin 12.4 (Vector Laboratories, Burlingame, CA), Alexa 633 0.07% (Thermo Fisher Scientific, Waltham, MA) and was adjusted to pH 7.3 with KOH. The standard Lucifer yellow pipette solution contained 10% H_2_O, 12.4 neurobiotin and 0.2% Lucifer yellow CH (Sigma-Aldrich, St. Louis, MO). For amphotericin recordings, cell-attached recording pipettes were filled with standard patch solution supplemented with freshly mixed amphotericin B (0.1 mM, Sigma). Liquid junction potentials were measured (9 to 11 mV), and the measured membrane potentials were corrected accordingly.

In the electrophysiological studies designed to record from identified light-responding mRGCs, we used retinas harvested from *Opn4^cre^::Ai96* embryos. To locate light-responsive ganglion cells, we first recorded the fluorescent response of the entire field of ganglion cells to ~5s epifluorescent stimulation (470/40 nm, 3 × 10^17^ − 3 × 10^19^ photons/cm^2^/s excitation, 525/50 nm emission). The locations of light-responsive fluorescent cells were marked on the screen monitor. The same area of the retina was then viewed with infrared DIC optics, and the patch pipettes were aligned with the cells showing light responses. Fluorescent and DIC images were collected using a CCD camera (Hamamatsu Digital Camera C4742) and Micro-Manager software (Open-image.com, San Francisco, CA).

### Intracellular calcium measurements

#### GCaMP calcium measurements

Epifluorescence from isolated retinas was imaged in an inverted Nikon (TE2000S) microscope (20x S-Fluor objective, NA 0.75). A Lambda LS 175-watt stand-alone xenon arc lamp and power supply (Sutter Instruments, Novato, CA) provided the light for stimulation. The incident light passed through an FITC/TRITC filter set (51004v2 - peak excitation wavelength ~480 nm; Chroma Technology Corp., Bellows Falls, VT). Stimulus-evoked epifluorescence was focussed onto a CCD camera (either a Hamamatsu C4742; 1024*1280 pixels or a Hamamatsu Orca Flash 4.0 LT; 2048×2048 pixels, Hamamatsu Corp., San Jose, CA). The field of view was ~ 430 μm × 340 μm (C4742) or ~ 650 μm × 650 μm (4.0 LT). Images were acquired using 4×4 binning and had a resulting resolution of 512 × 512 pixels. Metafluor software (Molecular Devices, Inc., San Jose, CA) controlled stimulation and data acquisition. The excitation light simultaneously activated GCaMP3 or GCaMP6 and stimulated melanopsin photopigment. Light stimuli were presented at 2 Hz and 3 Hz. Stimulation/acquisition periods were 530 ms (2 Hz) and 330 ms (3 Hz). The photon flux at the retina was routinely 4 × 10^17^ photons/cm^2^/s. Regions of interest encircling the somas of GCaMP3-fluorescent ganglion cells were set manually on images recorded from the best-focussed position. Fluorescence images from complete fields of view were recorded continuously. The summed responses from all pixels within each separate ROI were also viewed in real time and recorded to disc. Photobleaching of GCaMP3 was shown to be minimal based on the very small decrease in fluorescence in GCaMP-expressing cells lacking photopigment. Photoresponses could be repeatedly and reproducibly measured for periods of up to 60 min.

#### Calbryte-590 indicator dye

##### Exogenous calcium dye loading

We used Calbryte-590™ AM dye (573 nm Ex/ 588 nm Em; AAT Bioquest, Sunnyvale CA) to monitor intracellular calcium. This dye localizes to the cytoplasm and is 10-fold brighter than Rhod-2 AM (Zhao et al. 2019). A 5 mM stock solution of the dye in anhydrous DMSO was prepared just prior to use and diluted to a working solution of 20 µM by adding stock Calbryte-590™ AM solution to carbogen-bubbled ACSF containing Pluronic F-127 (0.04% f.c.). Isolated retinas were incubated in the ACSF/Calbryte-590™ solution for 2 hours in the dark with continuous carbogen circulation; they were then rinsed with fresh ACSF, mounted ganglion cell side down on a polylysine-coated (0.1 mg/mL) glass coverslip and placed in a heated (37°C) perfusion chamber. Calcium imaging was performed on the same optical rig used for the GCaMP experiments.

### Immunohistochemical labelling

For GCaMP3/melanopsin immunohistochemistry, the eye was removed, the cornea was perforated with a razor blade, and the entire eye was immediately immersed in 4% (w/v) paraformaldehyde (PFA) in PBS. In some experiments, the anterior part of the eye, including the cornea and lens, was first carefully removed before immersing the entire posterior eyecup in fixative. The eye was fixed in PFA in 0.1 M PBS for 30–90 min at RT. The eye was then stored overnight in 25% sucrose, 0.01% sodium azide in 0.1 M PBS at 4°C. The following day, the eyes were embedded in OCT and frozen on dry ice. Twenty-micron sections of the retina were cut perpendicular to the vitreal surface with a cryostat, mounted on Super-Frost Plus slides (Fisher Scientific, Pittsburgh, PA), and stored frozen at −20°C until immunostaining. ToPro3 (Life Technologies, Waltham, MA, Cat. # T3605) staining was used to mark nuclei. UF008 (1:500, Advanced Targeting Systems, San Diego, CA) and goat anti-GFP (1:500, Novus Biologicals, Littleton, CO) were used to label melanopsin and GCaMP3/GCaMP6, respectively.

### Drugs

FR900359 was isolated from the dried leaves of *Ardisia crenata* according to a previously published procedure (Schrage et al., 2015). YM-254890, tetrodotoxin, FPL65176 and L-cis-diltiazem were purchased from Wako Chemicals (Osaka, Japan), Calbiochem [MilliporeSigma] (Temecula, CA), Tocris Bio-Techne Corporation (Minneapolis, MN) and MilliporeSigma (Temecula, CA), respectively.

## Results

### Cells in foetal mouse retinas are light-responsive as detected by the fluorogenic calcium indicator dye Calbryte-590™

Melanopsin ganglion cells in foetal mouse pups differentiate before birth (Tartellin et al. 2003). In postnatal mouse retinas, mRGCs and perhaps other types of cells coupled to mRGCs are light-responsive, as demonstrated by stimulus-evoked increases in intracellular calcium levels (Sekaran et al. 2005). Photosensitive cells have not been reported in the retinas of embryonic mice or any other embryonic mammal. To determine whether such photosensitive cells exist, we searched for light responses by monitoring the activity of cells in embryonic mouse retinas after exogenously bulk-loading the retinas with the calcium-sensitive dye Calbryte-590™ AM (Zhao et al. 2019). After harvesting, the flat-mounted retinas were fluorescently imaged, immersed in Calbryte-590™ AM, mounted in a perfusion chamber, bathed with ACSF at 35°C, and stimulated with light of wavelength 480 nm (see Methods). Figure 1A shows an example of a Calbright-590™-loaded 340 μm × 430-μm region cropped from the full image (650 nm × 650 nm). This retina was harvested from an E17.5 mouse pup. The image was captured from the video recording at the onset of the light stimulus. Figure 1B shows an image of the same retina captured 2.5 s after stimulus onset. Figure 1C, which is a difference image (Fig. 1B minus Fig. 1A), accentuates the light-evoked increase in fluorescence that reflects the increase in intracellular calcium in the cells. The time courses of calcium increases in four of the cells, marked with circular regions of interest, are plotted in Fig. 1D. Peak changes in calcium centred approximately 2 s after the onset of the light. In total, we recorded Calbryte-590™ images in 2 to 3 animals taken from 3 different litters at age E17.5 and 2 litters at E18.5. We commonly detected more than 100 light-dependent calcium responses in each 430 mm × 340 mm region of the retina. Analysis of 4 of the brighter cells is shown in Figure 1.

**Figure 1.**
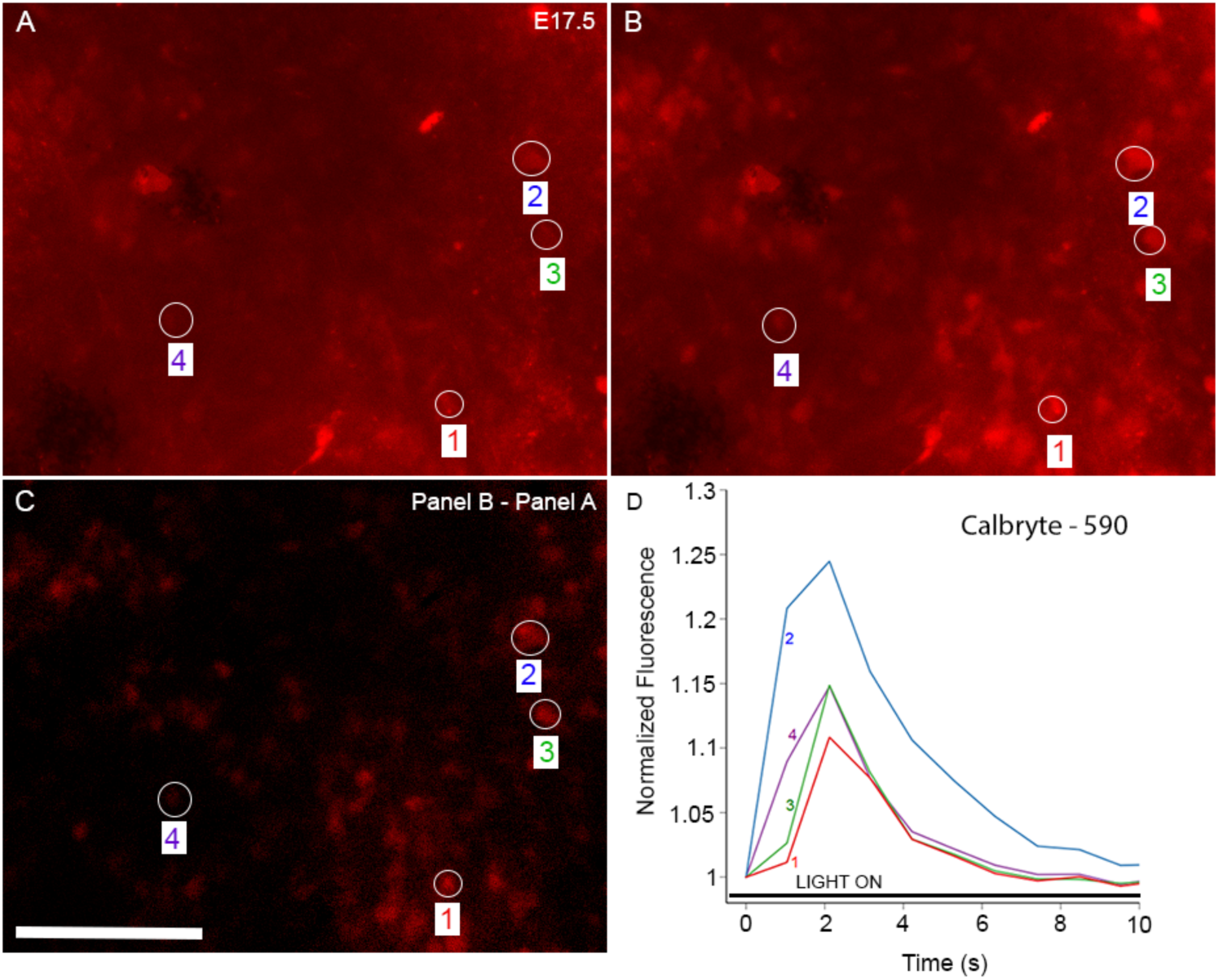
Cells in foetal mouse retinas are light-responsive. Cells in embryonic mouse retinas exhibit light-dependent increases in intracellular calcium. (A-C) Representative images of light-evoked fluorescence recorded from cells in an isolated, flat-mounted retina of an E17.5 mouse after loading of the retina with Calbryte-590 calcium-sensing dye. Images of a 340 × 430 mm region of the retina captured at T=0 s and at T = 2.5 s (A, B) are displayed. The colour-coded regions of interest (ROIs) show 4 cells in which light evoked a very discernible increase in fluorescence, indicating an increase in intracellular calcium. (C) Difference image of the two frames (T_2.5_ – T_0_) highlighting the regions in which increased dye fluorescence was observed after light onset. (D) Plots of the temporal response profiles of the cells marked by ROIs. The ordinate plots dF/F, where F is the baseline fluorescence immediately after the onset of the stimulus and prior to the onset of the response. Stimulus light of 480 nm wavelength was applied at 4 × 10^17^ photons/cm^2^/s. The scale bars in A, B and C indicate 50 μm.

Non-OPN4 opsins such as encephalopsin (OPN3) and neuropsin (OPN5) are expressed in the retina. Cells that express these opsins might also exhibit light-evoked Calbyte-590™ signals (Buhr et al., 2015, Kumbalasiri and Provencio, 2005). In mouse eyes, markers for neuropsin and encephalopsin appear after 10.5 days of gestation (Tarttellin et al. 2003). We tested foetal retinas from melanopsin knockout mice (*Opn4^cre^* homozygotes) and never observed a light-evoked increase in Calbright-590™ fluorescence in these retinas. These findings support the interpretation that the light-evoked calcium increases we observed in foetal retinas were evoked in OPN4 cells and that no detectable calcium elevations were observed in OPN3- or OPN5-expressing cells. It is noteworthy that light stimulation of OPN5 cells has been shown to modulate ocular dopamine levels in early postnatal mice (Nyugen et al. 2019).

In summary, the data presented here provide strong evidence that a subpopulation of cells within the foetal retina exhibits light-dependent calcium responses. However, we cannot rule out the possibility that the observed light responses reflect light-evoked activity from non-mRGCs as well as from mRGCs. For example, Calbryte-590™ AM dyes (and likely other AM indicator dyes) can indiscriminately load into cells of various types, including astrocytes (Tirschbirek et al. 2015). Thus, neurons or glial cells coupled to mRGCs could also exhibit light-evoked activation (Sekaran 2003; Reifler 2017). To address these ambiguities, we designed an experimental approach to selectively restrict the expression of calcium indicators to mRGCs.

### Melanopsin ganglion cells in foetal retinas are light-responsive as shown by the responses of the calcium indicator GCaMP3 expressed exclusively in mRGCs

We first asked whether the genetically encoded calcium indicator GCaMP3 is selectively expressed in mRGCs of mouse retina. We utilized the Ai38 mouse strain in which GCaMP3 is conditionally expressed. In the progeny of *Opn4^cre^*^/+^::*Ai38* mice, an allele that encodes the EGFP-based Ca^2+^ reporter GCaMP3 is transcribed in cells that express Cre recombinase (Zariwala et al. 2014). Because the *Opn4* gene is expressed relatively early in gestation, we reasoned that we could expect to observe conditional GCaMP3 expression well before birth and that it would be possible to detect light-evoked calcium signals that were restricted to melanopsin-expressing cells. Figure 2 A-C shows flat-mounted retinas in which melanopsin immunofluorescence co-localizes with EGFP immunoreactivity. This confirms that GCaMP3 is expressed in mRGCs. Of note, we also observed a population of GCaMP3-positive cells that showed little, if any, melanopsin immunoreactivity. The melanopsin-immunoreactive cells we did observe are likely to be predominantly mRGCs of the M1 subtype, which express high levels of melanopsin (Ecker et al. 2010). The other GCaMP3-EGFP-labelled cells may correspond to other mRGC subtypes that express melanopsin Cre but are known to express lower levels of melanopsin. In radial cryosections, an EGFP signal was evident in the inner plexiform layer and was strong in neurons with RGC morphology (Fig. 2D). These anatomical findings demonstrate that GCaMP3 is indeed localized to mRGCs. It should be noted that the images shown in Figure 1A-D were obtained from postnatal mice. Although CGaMP3 was well expressed in embryonic retinas, melanopsin antibody staining of prenatal retinas was not consistently detected. Therefore, based on the co-localization of antibody staining, it cannot be persuasively argued that the GCaMP3 cells in the foetal retinas co-expressed functional melanopsin protein. However, given that we were able to record light-evoked calcium increases in most of the embryonic GCaMP3-positive cells of the inner retina, there is little doubt that melanopsin protein was expressed in these foetal RGCs. The light sensitivity of mRGC-GCaMP3 embryonic neurons in retinae was next characterized.

**Figure 2.**
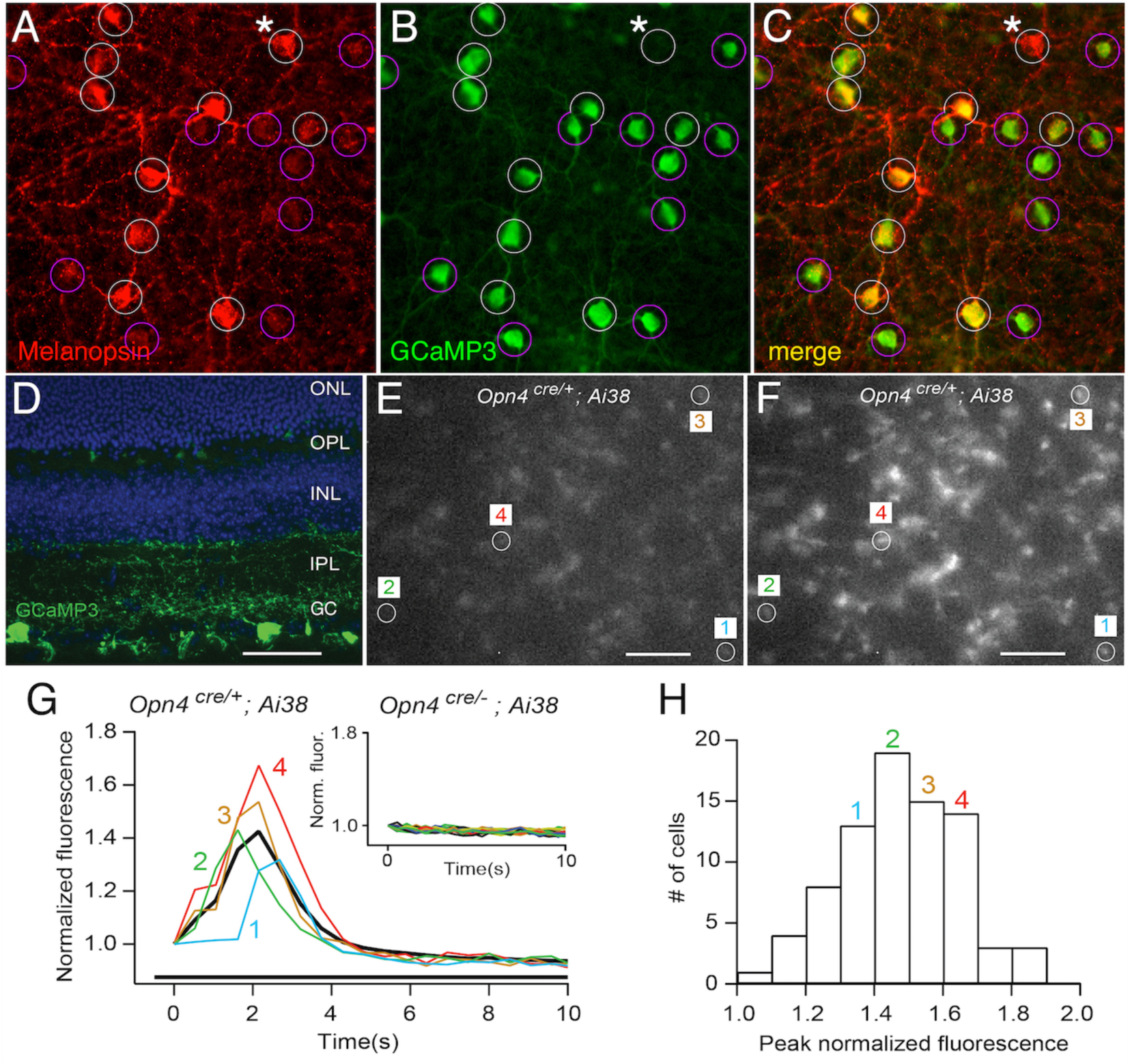
The genetically encoded intracellular calcium indicator GCaMP3 reveals light responses of melanopsin RGCs in embryonic retinas. Retinas from melanopsin knockout mice were unresponsive to light. GCaMP3-EGFP co-localizes with melanopsin and indicates light responsiveness of foetal mRGCs in *Opn4^cre/+^*::Ai38 retinas. (A) A flat-mount retina from an *Opn4^cre/+^::Ai38* postnatal day 7 (P7) pup showing its melanopsin immunoreactivity (red). (B) GCaMP3-EGFP-immunoreactive cells (green). (C) Merged image of (A) and (B). The white circles show co-labelled cells; the purple circles mark cells in which GCaMP3^+^ immunoreactivity is strong and melanopsin immunoreactivity is absent or very dim. White circles with an asterisk mark the occasional melanopsin+, GCaMP3-negative cells. These particular cells probably reflect failure of the GCaMP3 allele to recombine. (D) GCaMP3-EGFP immunoreactivity in a perpendicular cryosection of P30 retina. Immunoreactivity is observed in the ganglion cell layer (GC), the inner plexiform layer (IPL) and within a few structures in the outer plexiform layer (OPL). (E, F) Retinas isolated from embryonic pups were placed in a perfusion chamber and stimulated with 480 nm light. (E) GCaMP3 fluorescence at the onset of the light response in one representative E16.5 retina. (F) Fluorescence recorded at the peak of the light response in the same retina. Visual comparison of the images in (E) and (F) shows qualitatively brighter images in (F). (G) Time course of light-induced increases in df/F for 4 light-responsive cells (numbered 1 to 4 and colour-coded). The thick black trace plots the average response of 79 light-responsive cells in this experiment. These plots allow a more quantitative comparison of light responses in individual mRGCs. The scale bars indicate 50 μm. (G inset) Deletion of melanopsin (genotype *Opn4^cre/cre^::Ai38*) reveals, first, that melanopsin is required for light responses and, second, that the observed light-evoked increases in fluorescence do not result from unrecognized effects of light on GCaMP3-EGFP.

To search for light responses in foetal eyes, we mechanically isolated retinas from foetal pups, placed them in a perfusion chamber and stimulated them with light at 480 nm, the wavelength of light to which the melanopsin photopigment is most sensitive. Light stimulation resulted in [Ca^2+^]_i_ increases in GCaMP3-expressing neurons localized across the ganglion cell layer of the isolated retinas. Single-frame images of GCaMP3-filled RGCs show fluorescence in the retina of an E16.5 mouse before the onset of the stimulus (Fig. 2E) and during a light response (Fig. 2F). The complete time courses of changes in [Ca^2+^]_i_ in the 4 encircled cells (Fig. 2E) are shown in Fig. 2G. The black line in Fig. 2G shows the averaged response of all 79 cells recorded in this experiment. These records reveal that the time-to-peak following light onset ranged from 1.5 s to 4 s. The response kinetics were comparable to those reported in previous studies of postnatal mRGCs (Ecker et al. 2010, Hartwick et al. 2008, Sekaran et al. 2005). We also measured and plotted a cumulative histogram of peak response amplitudes (Fig. 2H). The responses of the individual cells (Fig. 2G) were taken from the 20^th^, 40^th^, 60^th^ and 80^th^ percentile groups (Fig. 2H). We discerned no correlation between peak amplitude and time course and no obvious partitioning of responses, suggesting that the temporal differences noted in different mRGC subtypes in postnatal mice are not evident at this age (Tu et al., 2005).

In total, we examined 1 to 4 littermates from 12 different embryonic litters. In *Opn4^cre/+^::Ai38* foetal pups, we found light-responsive cells in 6 of 6 litters. In *Opn4^cre/+^::Ai96* foetal pups, we observed light responses in 16 of 16 litters. Using our standard irradiance stimulus (3.8 × 10^16^ photons/cm^2^/s), we recorded up to 90 responsive cells in 430 μm × 340 μm regions of the retinas. The fact that as many as 90% of the GCaMP3-positive cells exhibited light responses argues against the possibility that artefactual Cre expression may have occurred in non-melanopsin-producing cells (Delwig et al., 2016; Ecker et al., 2010). We were able to find detectable [Ca^2+^]_i_ elevations with stimulus irradiances as low as 3 × 10^16^ photons/cm^2^/s. At stimulus irradiances below that level, the signal-to-noise ratios were too low for reliable detection.

No light responses were observed in control experiments performed in melanopsin knockout retinas. *Opn4^cre/+^::Ai38* mice crossed with *Opn4^cre/cre^* mice produce some *Opn4^cre/cre^::Ai38* pups. The mRGCs in these pups express GCaMP3 but not melanopsin. We found no light-responsive cells in these foetal pups (Fig. 2G, inset). The experiments with knockout retinas also serve to rule out the possibility that the light-evoked increase in GCaMP3 fluorescence in melanopsin-expressing cells is an artefact unrelated to melanopsin-mediated calcium influx. Additionally, the recordings in knockout cells reveal that bleaching of GCaMP3 is minimal under our experimental conditions. Specifically, the fluorescence amplitude 10 seconds after light onset was 94 ± 3% of the initial amplitude (Fig. 2G, inset).

In summary, these results convincingly demonstrate that mRGCs in foetal mouse retinas are responsive to light. In addition, the Calbryte590™ measurements in OPN4 knockout mice suggest that any photo-responses generated by other opsins, such as OPN3 and OPN5, were below the limit of detection.

### Antagonists of the G_q/11_ G-protein family attenuate mRGC light responses in foetal retina

Light-activated opsins in invertebrate eyes signal via coupling to members of the G_q/11_ family of G-proteins (Katz B & Minke B., 2009). Similarly, light-activated melanopsin in mRGCs has been postulated to signal via a comparable G_q/11_ pathway (Panda S et al., 2005; Hughes et al., 2015; Graham DM et al., 2008; Jiang et al., 2018). However, a recent study reported that deletion of G_q/11_ G-proteins does not abolish light responses in postnatal mRGCs (Chew et al., 2014). We tested whether G _q/11_ signalling plays an essential role in embryonic mRGC light responses. To do this, we examined the effects of two different cell- permeable antagonists, FR900359 (Schrage et al., 2015) and YM-254890 (Kawasaki et al., 2003; Nishimura et al., 2010), on light responses. These two compounds inhibit activation of the Gnaq, Gna11 and Gna14 alpha subunits of G_q/11_-G proteins. For these experiments, we used *Ai96* mice that express the calcium indicator GCaMP6. GCaMP6 has superior sensitivity and a better signal/noise ratio than GCaMP3 (Chen et al., 2013). The methods of breeding and fluorescence recording were otherwise identical to those used with the *Ai38* animals in the previous experiments. Recordings were made in *Opn4^cre/+^::Ai96* retinas harvested from E15.5 and E16.5 foetal pups. The retinas were placed in control medium (ACSF), and light responses were recorded from mRGCs before and after the addition of YM-254890 at 200 nM or FR900359 at 1 μM. Figure 3A shows an example of a recording from one retina; the light responses, quantified as the increase in light-induced fluorescence, dF/F, were stable over a 40-min period in ACSF but were progressively attenuated during the 30-min period following the addition of YM-254890. Four representative light responses on an expanded time scale are shown (Fig. 3B). We found no evidence of reversibility following washout. Poor reversibility has been reported previously (Yasui et al., 2008). The time courses of the mean light responses from multiple mRGCs in each retina are shown in Fig. 3C and D. Both YM-254890 (E15.5 retina, 32 mRGCs) and FR900359 (E16.5 retina, 43 mRGCs) severely attenuated light responses after 30 min. To rule out the possibility that the inhibitors merely killed the mRGCs or quenched the fluorescence, we applied high potassium (50 mM) at the end of the inhibitor infusion. Figure 3D shows an example in which potassium substitution was able to induce a dF/F larger than that originally induced by light.

**Figure 3.**
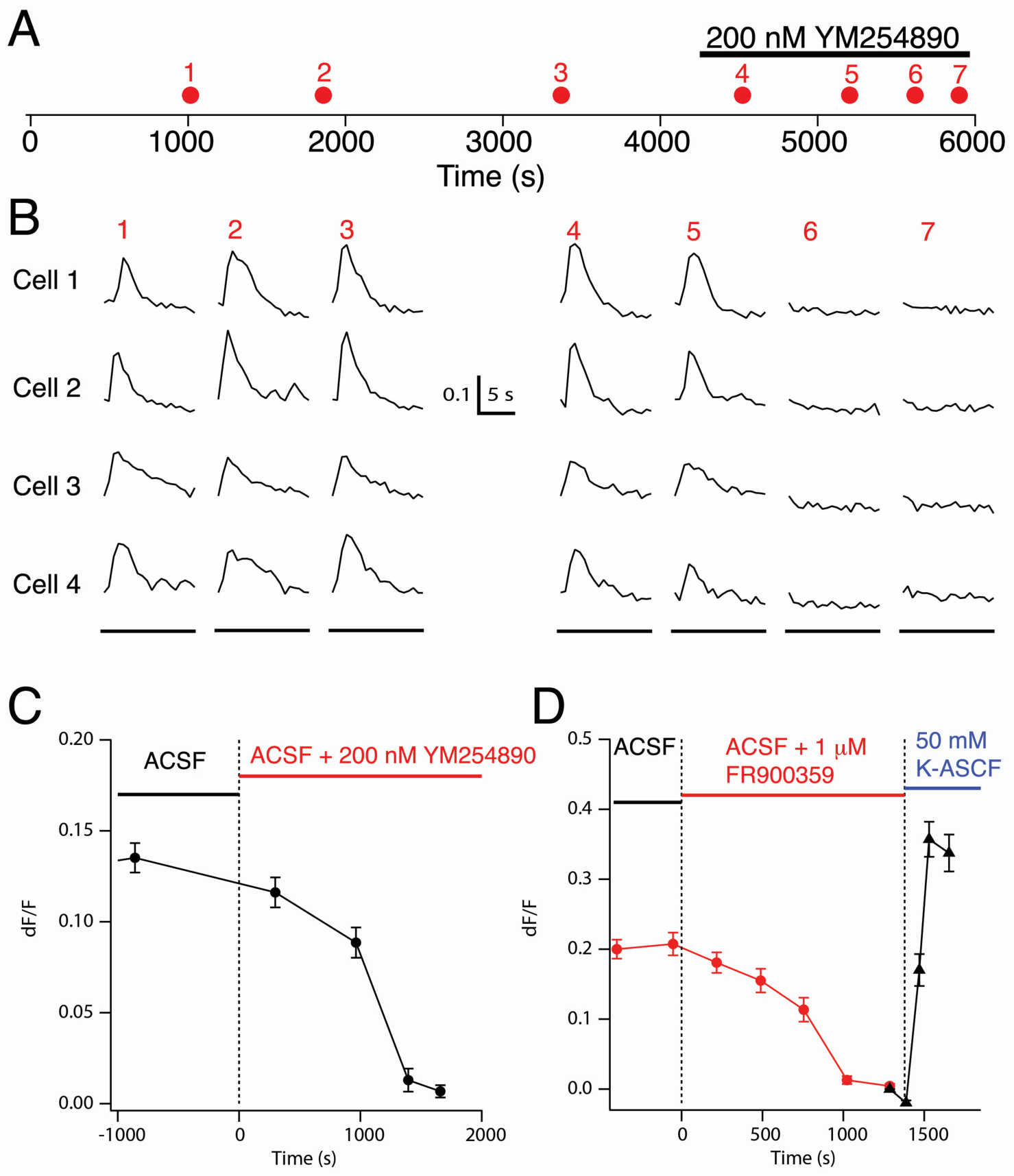
Antagonists that block the functions of the Gnaq, Gna11 and Gna14 alpha subunits of G_q/11_-G proteins of the G_q/11_ family severely attenuate the light responses of mRGCs in foetal retinas. The effects of YM-254890 (200 nM) and FR900359 (1 μM), two different antagonists of Gnaq, Gna11 and Gna14, on light-evoked calcium elevations were tested in 4 explanted E15.5 and 3 explanted E16.5 OPN4cre::Ai96 retinas. (A) Time course of the experiment in an E16.5 retina showing 12-s light stimuli (red dots, 1-7) over the course of 6000 s. Responses to light stimuli 1-3 were recorded during 3300 s in control ACSF, and responses to light stimuli 4-7 were recorded after application of YM-254890. (B) Selected light responses from 4 mRGCs before and during perfusion with YM-254890. The scale bar shows df/F of 0.1. (C) Mean response amplitudes observed in 32 simultaneously recorded mRGCs in the same E15.5 retina analysed in (B). (D) Mean response amplitudes observed in 43 simultaneously recorded mRGCs in an E16.5 retina perfused with medium containing 1 mM FR900359 (from ~400 to 1800 s). When dF/F responses were close to zero at 1400 seconds, the retina was perfused with 50 mM K-ACSF in which 47.5 mM NaCl was substituted for 47.5 mM KCl. The inclusion of KCl in the ACSF increased the calcium fluorescent signals to a level brighter than that induced by light before perfusion with FR9000359. All of the cells plotted in (C) and (D) had an initial criterion light response dF/F > 0.05. The error bars indicate the SEM.

Prolonged incubation in 200 nM YM-254890 reduced the average dF/F of the light responses to 7% ± 11% of that of the control responses (n=93). Prolonged incubation in 1 μM FR900359 reduced the average dF/F of the light responses to 4% ± 18% that of the control responses (n=219). Comparing light responses from individual cells prior to and during drug application, we found highly significant drug effects across the ensemble of responsive cells. According to the Wilcoxon Signed Rank Sum Test for paired data, the two-tailed P = 2.01948e^−28^ for YM and was even smaller for FR.

The use of two independent Gnaq/11/14 inhibitors diminishes the possibility that off-target effects of either FR900359 or YM-254890 are responsible for the observed suppression of light responsiveness. These results implicate G_q/11_ signalling in the generation of light responses in embryonic mRGCs. One caveat is that the averaged data shown in Figure 3 may not reflect the presence of subpopulations of mRGCs that might utilize non-G_q/11_ pathways.

### No subgroups of G _q/11_-independent mRGCs in embryonic retinas are detected in population data

Jiang et al. (2018) reported that in adult mice most classes of mRGCs employ a G_q/11_ cascade for transduction. However, they also reported that M4- type mRGCs utilize a second alternative transduction cascade involving a cyclic nucleotide as the second messenger and HCN channels. Jiang et al. (2018) reported that this pathway very likely does not use G _q/11_ family members or TRP6/7 ion channels. In support of this conclusion, they showed that M4 mRGCs responded to light normally in cells devoid of *Gnaq*, *Gna11* and Gna14 as generated in Gnaqfl/fl;Gna11^−/−^;Gna14^−/−^mouse eyes transfected with Cre-GFP via adeno-associated virus serotype 2 (AAV2-CMV-Cre-GFP).

We hypothesized that if a group of mRGCs in foetal retinas also does not use G _q/11_ signalling, then a subpopulation of GCaMP6-expressing mRGCs should be resistant to the action of YM and FR. Figure 4, below, shows cumulative data from 93 mRGCS tested in YM-254890 and 219 mRGCs tested in FR900359. Our data indicate that only approximately 1% of the cells might be resistant to these drugs. Thus, our experiments provide no supportive evidence for the existence of a detectable subpopulation of foetal mRGCs that utilize a non-G _q/11_ transduction pathway.

**Figure 4.**
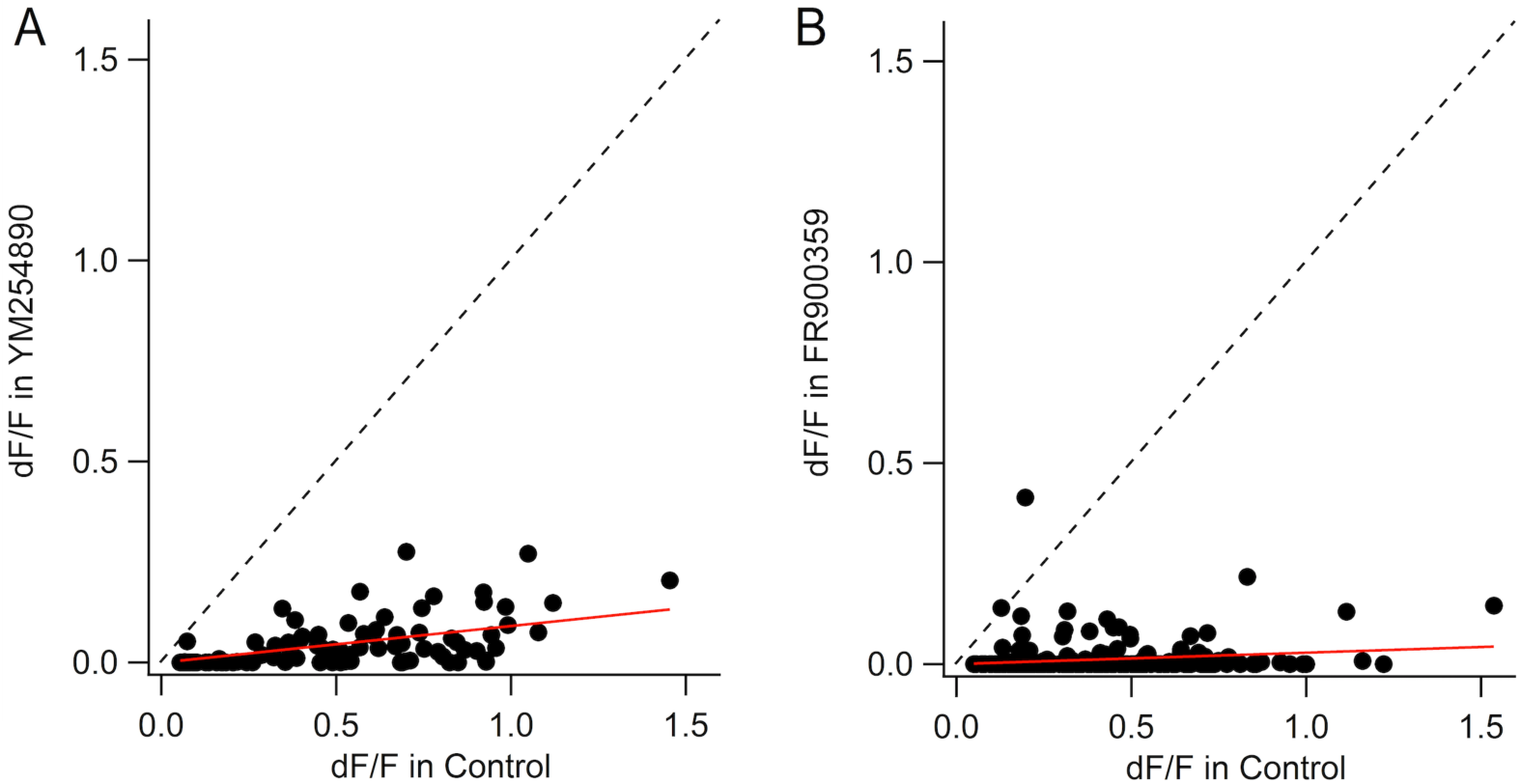
Ensemble data confirming that YM254890 and FR900359 uniformly attenuate light responses in virtually all embryonic mRGCs. (A) Combined light-evoked dF/F’s of 93 cells from E15.5 and E16.5 *OPN4cre::Ai96* retinas. All cells had a light-evoked dF/F > 0.05 in control conditions. The black dots indicate the lowest dF/F after 1500-2500 s incubation in 200 nM YM-254890 versus control dF/F for each cell. Prolonged incubation in 200 nM YM-254890 reduced the average dF/F of the light responses to 7% ± 11% that of the control responses (n=93). The data are best fitted by the linear regression y = 0.09x (solid red line). The dashed line indicates the predicted location of the dots if YM-254890 had no effect. (B) Combined light-evoked dF/F’s of 219 cells from 3 embryonic day 16.5 retinas. All cells exhibited dF/F > 0.05 under control conditions. The black dots indicate the lowest dF/F after a 1300-3100 s incubation in 1 mM FR900359 versus control dF/F for each cell. Prolonged incubation in 1 mM FR900359 reduced the average dF/F of the light responses to 4% ± 18% that of the control responses (n=219). The data are best fitted by the linear regression y = 0.03x (solid red line). These results are highly significant (Wilcoxon Signed Rank Test for paired data [two-tailed P = 2.01948e^−028^ for YM and even smaller for FR]).

### TTX-sensitive sodium currents amplify mRGC light responses in embryonic retinas

Based on electrophysiological recording and calcium measurements from neonatal mRGCs harvested from rat retinas, Hartwick et al. (2007) proposed a transduction pathway in which melanopsin-driven G-protein activation leads to the opening of cationic TRPC6/7 channels. The resulting inward cation flux subsequently depolarizes the membrane, leading to activation of TTX-sensitive sodium action potentials and then to activation of L-type calcium channels, which, in turn, increases calcium ion influx. We sought to determine whether TTX-sensitive sodium and L-type calcium currents support light responses in embryonic mRGCs. The likelihood of finding a role for TTX-sensitive currents in embryonic mRGC responses is enhanced by previous work demonstrating that a majority (78%) of randomly selected RGCs in E15 mouse retinas expressed TTX-sensitive sodium currents (Rorig and Grantyn, 1994).

We found that TTX (1 µM) reversibly diminished the average light response amplitudes of virtually all the foetal GCaMP6-expressing mRGCs we tested. Figure 5 shows the responses recorded from 177 cells in 4 E16.5 retinas. Each solid data point (blue) denotes the light response amplitude in TTX (ordinate) as a function of the amplitude in control ACSF prior to TTX (abscissa). The open circles show the responses of the same cells after washout of TTX. We found that TTX effectively reduced the light responses to below the detectable threshold in 21% of the cells (dF/F < 0.05). The aggregate differences between responses in TTX versus washout are highly significant (Wilcoxon Signed Rank Test for paired data, two-tailed P = 1.01e^−46^). Of note, these data show that while TTX reversibly diminished the light responses in virtually all of the tested mRGCs, the effects of TTX varied across the sample of cells. This variability is at odds with Hartwick et al. (2007), who reported that TTX abolished responses nearly completely in all mRGCs (n = 8). The observed difference in the efficacy of TTX could reflect our sampling of a larger population of mRGCs. Alternatively, these findings might reflect cell-specific differences in the maturation of INa(V) channels during embryonic and postnatal development. Nonetheless, we conclude that these TTX-sensitive channels do serve to amplify the light-evoked responses in a plurality of embryonic mRGCs.

**Figure 5.**
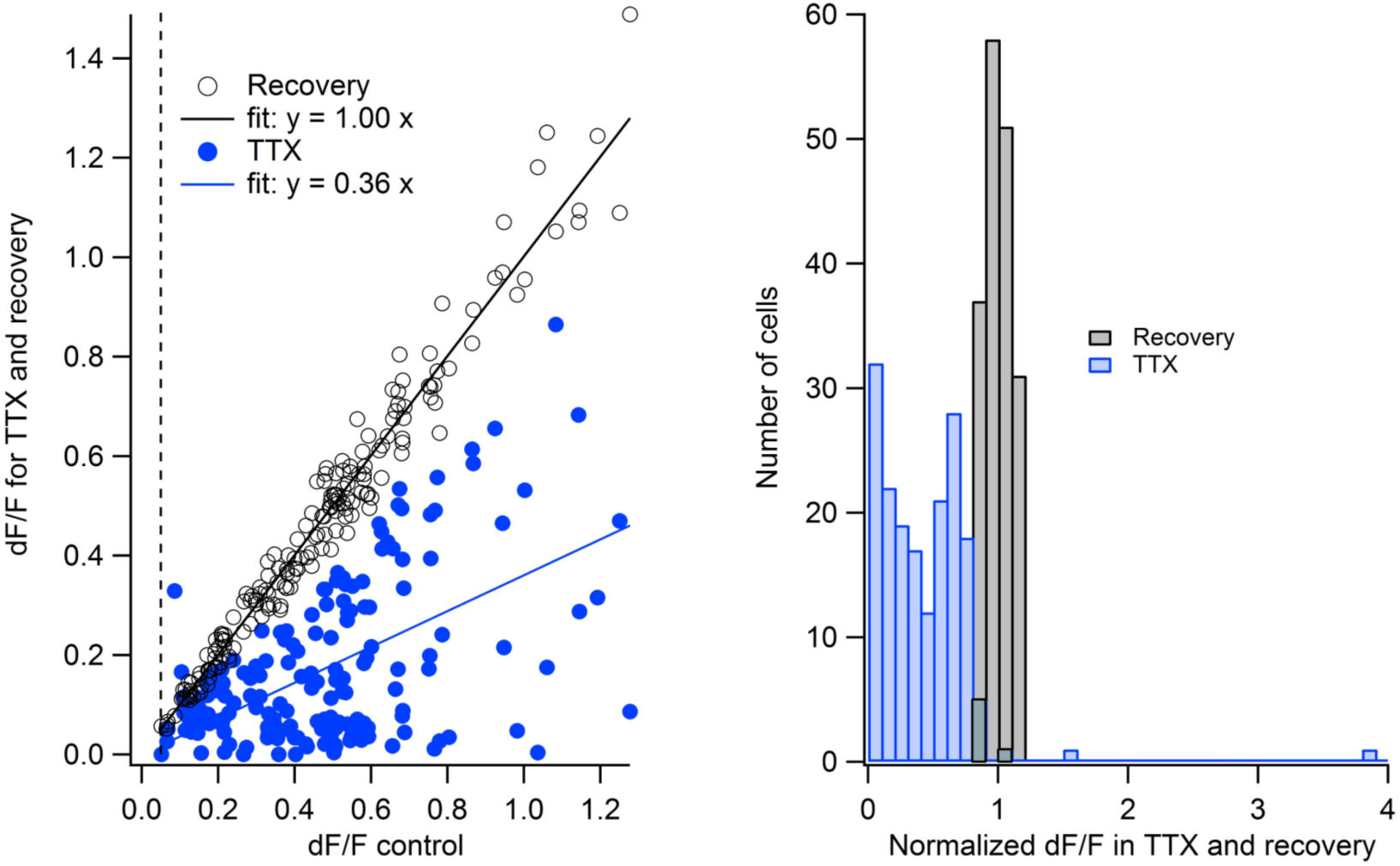
TTX variably but reversibly reduces the amplitude of light responses across the population of embryonic mRGCs. (A) We measured the action of TTX on the fluorescent light responses of GCaMP6-expressing mRGCs in *OPN4cre::Ai96* embryonic retinas. Combined data from 4 E16.5 retinas are shown. The criteria for selecting each cell were that dF/F in control ACSF was > 0.05 and that the response amplitude after TTX washout recovered to between 80% and 120% of the control value. The open circles plot the recovery dF/F versus control dF/F for each of the 177 light-responsive mRGCs. The solid blue dots plot dF/F in 1 μM TTX versus control dF/F for each cell. The black line (y = 1.00 x) indicates the best-fitting linear regression line through the recovery data and the origin 0,0. The blue line (y = 0.36 x) is the best-fitting linear regression line for the TTX data. The differences between the results obtained during TTX treatment and under control conditions are highly significant (Wilcoxon Signed Rank Test for paired data, two-tailed P = 1.01e^−46^). (B) Histogram of the data shown in (A). The blue bars plot dF/F in TTX divided by the dF/F under control conditions, and the grey bars plot dF/F in recovery divided by the dF/F under control conditions.

### Modifiers of L-type voltage-gated calcium channels modulate the light responses of mRGCs in E16.5 mouse retinas

Previously, Hartwick et al. (2007) showed that L-type voltage-gated calcium channels account for most of the calcium influx that occurs following light stimulation of enzymatically isolated postnatal rat mRGCs. While there are no reports of L-type calcium channels specifically in mRGCs of embryonic retinas, earlier studies suggested that L-type calcium channels are expressed in unspecified, randomly selected RGCs in E15 mouse retinas (Rorig and Grantyn, 1994). Consistent with this finding, expression of two L-type channel subunits, Cacna1a and Cacna1b, in the eyes of E15.5 mouse pups has been reported (Allen Brain Atlas, Cacna1b - RP_090707_03_G11 - sagittal). We tested pharmacologically for evidence that L-type calcium channels contribute to the light responses of GCaMP3-expressing mRGCs. In one set of experiments, we found that L-*cis*-diltiazem, a blocker of L-type calcium channels (Catterall et al., 2005), significantly diminished light responses in embryonic retinas (n=3). Figure 6 shows the results obtained from 31 cells that were recorded simultaneously in one retina. Panel A shows examples of reversible L-cis-diltiazem-induced suppression of light responses in 5 mRGCs. Panel B plots the responses from all mRGCs in which the light response peak amplitude returned to a value between 80 and 120% of that observed under control conditions upon washout. The blue dots plot the responses in L-cis-diltiazem, and the black dots plot the amplitude of the responses following washout. The five circled data points correspond to the light responses shown in Fig 6A. The effect of L-cis-diltiazem is highly significant according to the Wilcoxon Signed Rank Test for paired data (two-tailed P = 9.31e^−10^).

**Figure 6.**
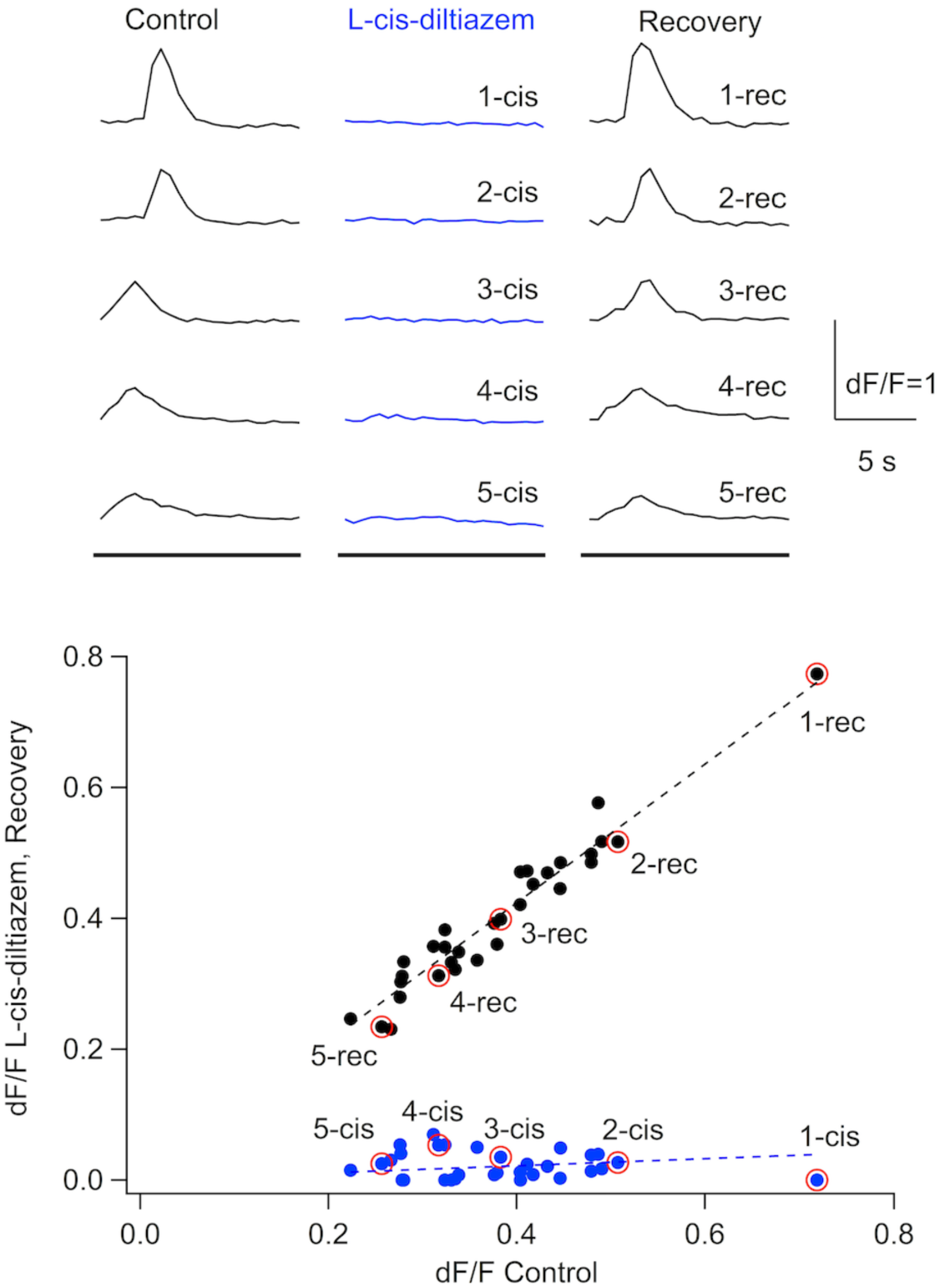
The L-type calcium channel antagonist L-cis-diltiazem (LCD) reversibly attenuates light-dependent increases in calcium in E16.5 mRGCs. (A) dF/F versus time in control ACSF, in LCD and after return to ACSF. Plots show representative recordings for 5 of 48 light-responsive cells from a single E16.5 *OPN4cre::Ai96* retina. The bars at the bottom indicate the timing of the light stimuli. (B) Plots of dF/F in LCD and recovery versus dF/F in control for 31 of 48 light-responsive cells in one retina in which the post-washout values recovered to between 80 and 120% of the control values. The dotted lines show the best linear regression that fits the data (y = 1.06x and y = 0.05x for recovery and LCD, respectively). The data from the 5 cells shown in (A) are shown as red circles. The Wilcoxon Rank Test for paired data yields a two-tailed P-value of 9.31323e^−10^, indicating that L-cis-diltiazem at 25 μM highly significantly diminishes light responses in mRGCs.

In a complementary set of experiments, we found that FPL64176, an activator of L-type calcium channels (Zheng et al., 1990), increased light-evoked dF/Fs in embryonic mRGCs. In previous studies, FPL64176 was shown to enhance the activity of retinal L-type calcium channels. Specifically, FPL64176 enhanced L-type calcium-mediated evoked release of the neurotransmitter glycine from amacrine cells (Bieda & Copenhagen, 2004). In addition, FPL64176 potentiated spontaneous waves of activity that occur in neonatal retinas (Singer et al., 2001). Figure 7 plots the peak light-evoked fluorescent responses observed before and during exposure of explanted retinas to FPL64176 (2 μM). The data include the responses from 147 GCaMP3-expressing mRGCs from 2 different E16.5 retinas. The responses observed during treatment with FPL64167 are plotted against the responses observed under control conditions. On average, the df/F values in FPL64176 were 0.22 ± 0.02 (SEM) larger than dF/F in control. The blue line shows the best linear regression fit to the FPL64176 data (y = 1.51x). The black line shows the values that would be expected if FPL64176 had no effect on the light responses. The Wilcoxon Rank Sum Test for paired data yielded a two-tailed P value of 1.172e^−32^. Together, the L-cis-diltiazem and FPL-64176 findings strongly support a role for L-type calcium channels in the generation of the light responses, similar to the transduction role proposed by Hartwick et al. (2007) for postnatal mRGCs.

**Figure 7.**
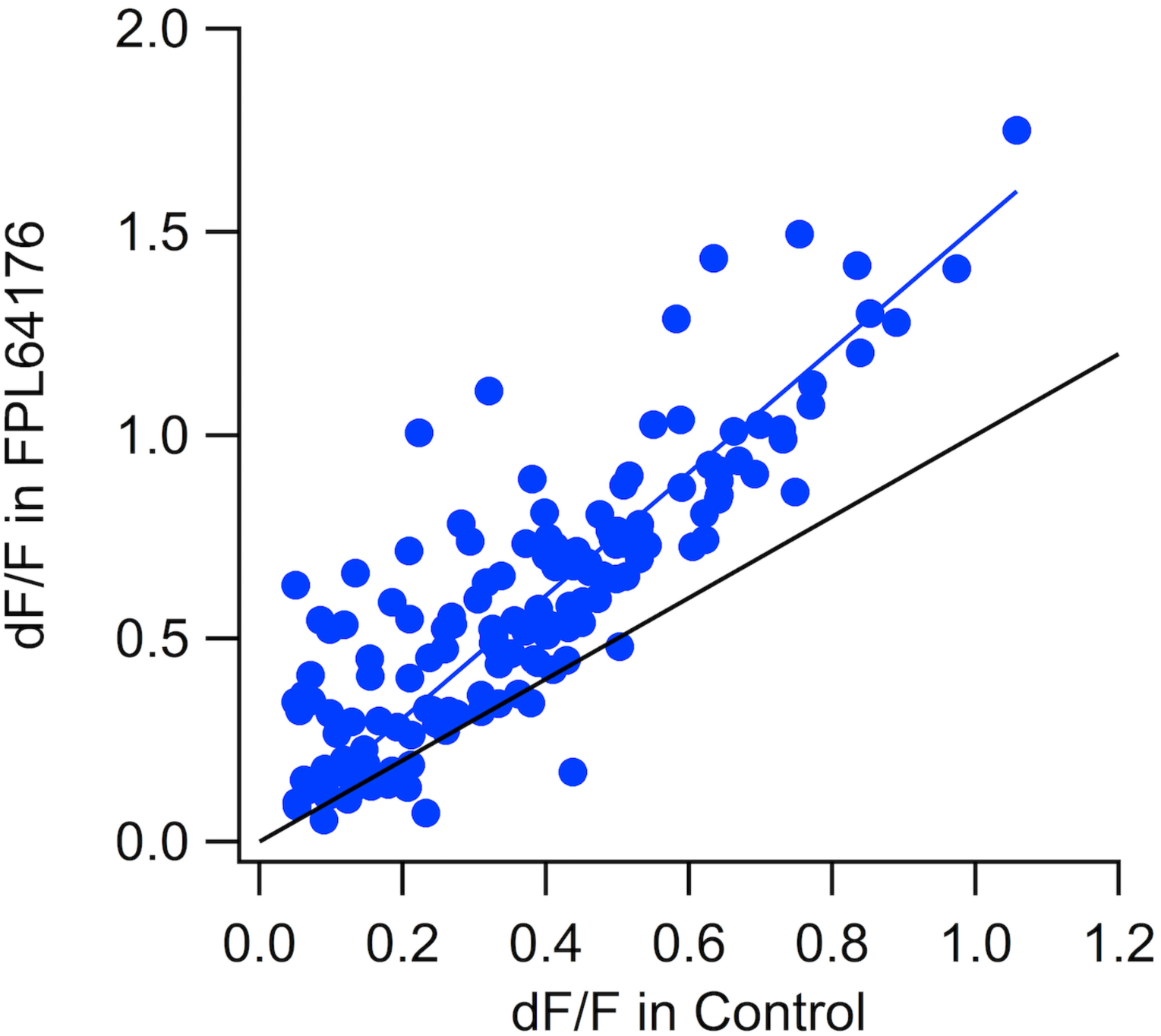
The L-type calcium channel agonist FPL64176 increases light-evoked dF/Fs recorded from embryonic mRGCs. A) dF/F in 2 mM FPL64176 versus dF/F in control for 147 mRGCs combined from 2 E16.5 *OPN4cre::Ai96* retinas. The blue line represents the best linear regression fit: y = 1.51x. The black line represents the parity of response in control and FPL64176. The Wilcoxon Rank test for paired data yielded a two-tailed P-value of 1.17193e^−32^, indicating that dF/F is very significantly larger during exposure of the mRGCs to FPL64176 than under control conditions. No recovery data are shown, as FPL washout is known to be very poor (Bieda and Copenhagen, 2004).

### Voltage-activated sodium currents exist ubiquitously in mRGCs from at least E16.5

Having found that the INa (V) antagonist TTX reduces and sometimes completely abolishes light responses in embryonic mRGCs, we sought to characterize these sodium currents electrophysiologically. Using patch pipettes, we recorded from mRGCs genetically marked with either GCaMP6, tdTomato or EGFP derived from *Opn4*^cre^::Ai96, *Opn4^cre^*::Ai14, or *Opn4^EGFP^* mouse strains, respectively. Patch pipettes were directed towards the fluorescently marked cells in isolated flat-mounted retinas. In the GCaMP-labelled retinas (*Opn4^cre^::Ai96*), we recorded from only those RGCs that displayed light-evoked calcium responses as reported by GCaMP6. An example of a voltage clamp recording from an mRGC in E18.5 retina is shown in Fig. 8. Panels A-C show current responses to a series of sequentially larger 10 mV voltage steps initiated at –70 mV (see Panel E). Brief, transient inward currents began to occur at – 40 mV and became faster with successive depolarizing pulses (Panel A). Perfusion with TTX (1 μM) reversibly blocked the inward currents (Panels B and C). The evoked TTX-sensitive currents (Panel D) were isolated by subtracting the response in Panel B from Panel A. The voltage dependence of the TTX-sensitive currents is plotted in Panel F. The peak current amplitude (black dots) exhibited a sharp onset at −40 mV, reached a maximum at −30 mV and then declined with further depolarizing steps. The kinetics and voltage dependency of the TTX currents shown here closely resemble those of the canonical INa(V) currents observed in excitable muscle and nerve cells across multiple phyla (Johnston & Wu, 1995).

**Figure 8.**
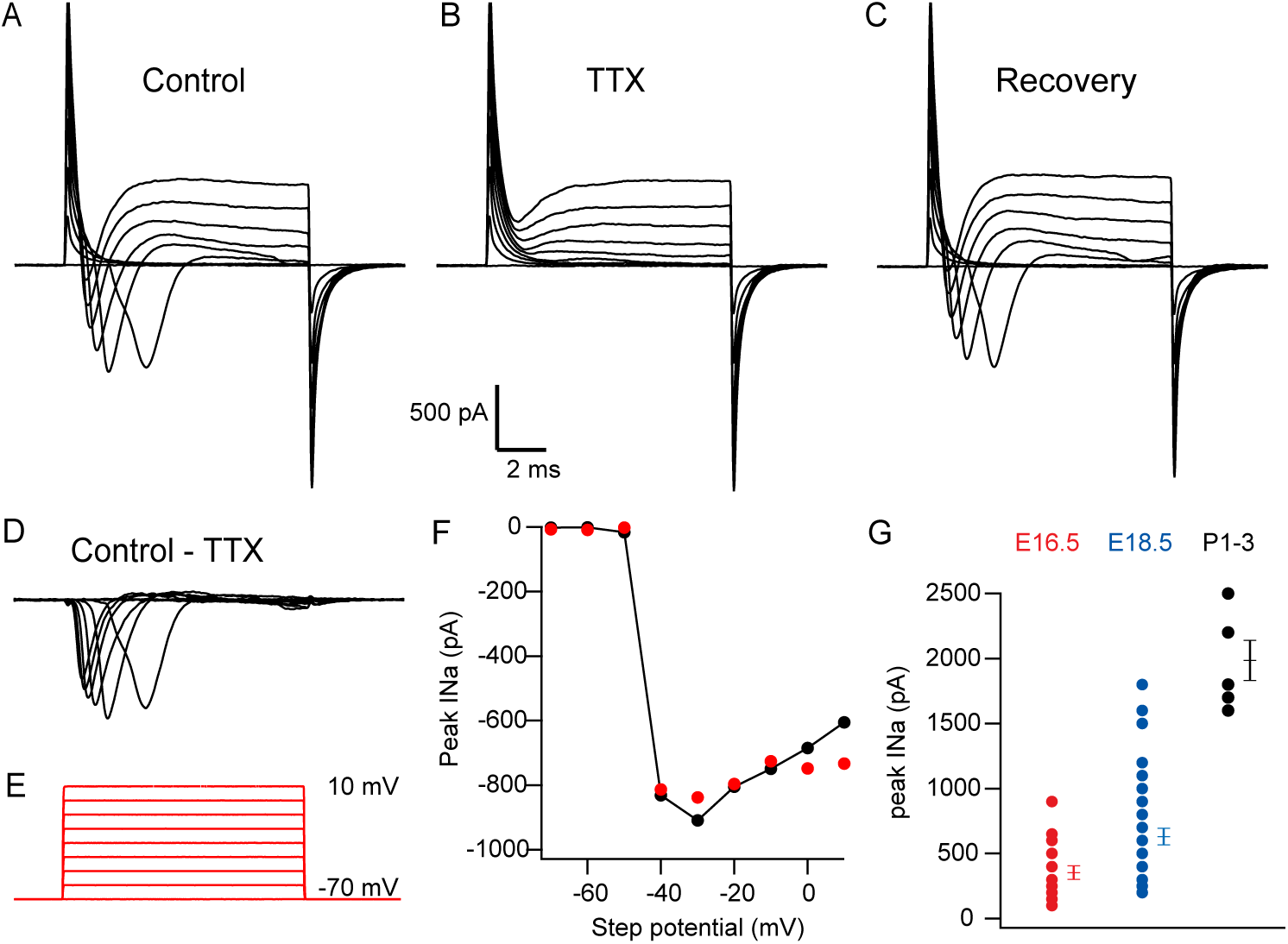
Patch pipette recordings reveal that TTX-sensitive sodium currents in mRGCs increase in amplitude during maturation from E16.5 to P1-3. A-F show examples of TTX-sensitive currents recorded from an mRGC in an E18.5 retina (*OPN4cre::Ai96*) under voltage clamp conditions. A) Currents evoked under control conditions with the voltage clamp protocol shown in E. B) Currents evoked after several minutes in 1 μm TTX. C) Currents evoked after washout of TTX. D) Difference current, Control - TTX. E) Voltage clamp step protocol. F) The black dots indicate the maximum inward current for each trace from D versus the step potential (measurement of INa(V). To survey the amplitudes of peak Ina(V) across multiple cells in retinas of 3 different ages without applying TTX, we used an analytical method in which the final current sample of the voltage step in A was subtracted from the lowest current amplitude across the entire voltage step. These values, plotted as red dots, yield a good estimate of peak INa(V), as shown by comparison with the results obtained in TTX (black dots). G) Peak sodium currents in mRGCs during embryonic maturation. The dots indicate peak INa(V) estimates for 10, 35 and 7 mRGCs recorded from E16.5, E18.5 and P1-3 retinas, respectively. The vertical bars indicate the average ± standard error for each age (320 ± 73, 674 ± 72 and 1986 ± 153 pA, respectively). The data include whole-cell and perforated recordings and were not corrected for series resistance. In summary, these findings demonstrate that voltage-activated sodium currents are evident in mRGCs at least 4 days before birth and that the magnitude of these currents increases with age.

### The mean amplitudes of INa (V) currents increase substantially from E16.5 to P1-3

INa (V) currents were detected in virtually all of the mRGCs recorded in 10 E16.5, 35 E18.5 and 7 P1-3 retinas. The peak INa (V) currents (not all data were analysed and included) are plotted in Fig. 8 (Panel G). The mean peak currents in the aforementioned groups of retinas were 320, 674 and 1986 pA, respectively. Comparable developmentally linked increases in the amplitude of INa(V) currents have been reported in random samplings of RGCs in mouse and rat retinas (Rorig and Grantyn, 1994; Schmid and Guenther, 1996). Our findings confirm that functional voltage-dependent sodium channels are expressed in mRGCs as early as E16.5 and further reveal an increase in the size of these currents increase during embryonic maturation. It is evident that the peak INa (V) varied across the sampled population at each age. We were unable to reliably and systematically determine whether the variability reflected the presence of cells of different sizes or whether the expression of INa (V) channels per unit of cell membrane varied. Of note, Schmid & Guenther (1996) divided whole-cell sodium current amplitude by cell capacitance and found that INa (V) current density increased sizably from E18 to P6.

### Electrophysiological recordings demonstrate that irradiance with light as dim as 3 × 10^12^ photons/cm^2^/s evoke responses in individual foetal mRGCs

Our previous work demonstrated that the ambient light in the mouse housing room was bright enough to trigger subsequent vascular development via stimulation of melanopsin in the eyes of foetal pups (Rao et al. 2013). Specifically, normal regression of the hyaloid blood vessels was retarded in foetal pups lacking expression of melanopsin (*OPN4^−/−^)* and in WT pups whose mothers were dark-reared from 16.5 days post-conception to the time of the pups’ birth. The effects of ambient light were not transmitted from the pregnant female to the pups but were a direct effect of ambient light on the foetal pups themselves. This was shown by the fact that environmental light still contributed to normal ocular vessel development when the pregnant female was enucleated or when melanopsin was selectively deleted in the pregnant mothers but not in the foetal pups. Thus, the light that struck the foetal eyes resulted from ambient light transmission through the abdominal walls of the dams. Direct measurements of the attenuation of the intensity of blue light across the dams’ skin and muscle layers support the notion that if mRGCs in embryonic retinas are as sensitive to light as mRGCs in postnatal retinas, the embryonic mRGCs could easily be activated by low-intensity environmental light at floor level in the mouse housing room (5.6 × 10^13^ photons/cm^2^/s, Rao et al. 2013). However, the actual photosensitivity of embryonic mRGCs is unknown. In the GCaMP experiments described above (Figs 2–7), we were not able to measure mRGC sensitivity because the minimum stimulus intensity needed to interrogate GCaMPs strongly activated melanopsin as well. To circumvent this experimental obstacle, we turned to a non-optical, electrophysiological approach using patch pipette recordings from individual mRGCs.

We recorded light-evoked responses from fluorescently marked mRGCs in foetal retina preparations. We found a total of 15 mRGCs that exhibited detectable light responses. Figure 9 shows light responses recorded from an mRGC in the retina of an E18.5 pup. Action potentials were observed at an irradiance of 2.9 × 10^12^ photons/cm^2^/s (Fig. 9D4). The onset of spiking displayed a latency of 21 s. The responses to steps of light at higher irradiances (5.8 × 10^14^, 1.5 × 10^14^ and 1.35 × 10^13^ photons/cm^2^/s) are plotted in Fig. 9D1, D2, and D3, respectively. Similar to the mRGC responses recorded in neonatal and adult retinas, the response latencies decreased as the irradiance increased (Tu et al. 2005, Wong KW, 2012). In our sample of cells, one mRGC had a threshold of 1.6 × 10^12^ photons/cm^2^/s. Seven of the 15 mRGCs had thresholds between 6.2 × 10^12^ and 3.5 × 10^13^ photons/cm^2^/s, and six of the 15 light responders had thresholds between 1.5 × 10^14^ and 2.4 × 10^14^ photons/cm^2^/s. It should be noted that all of the GCaMP6-expressing mRGCs recorded electrophysiologically were initially identified by their responsiveness to light at 3.10^17^ – 3.10^19^ photons/cm^2^/s (Fig. 9A,B).

**Figure 9.**
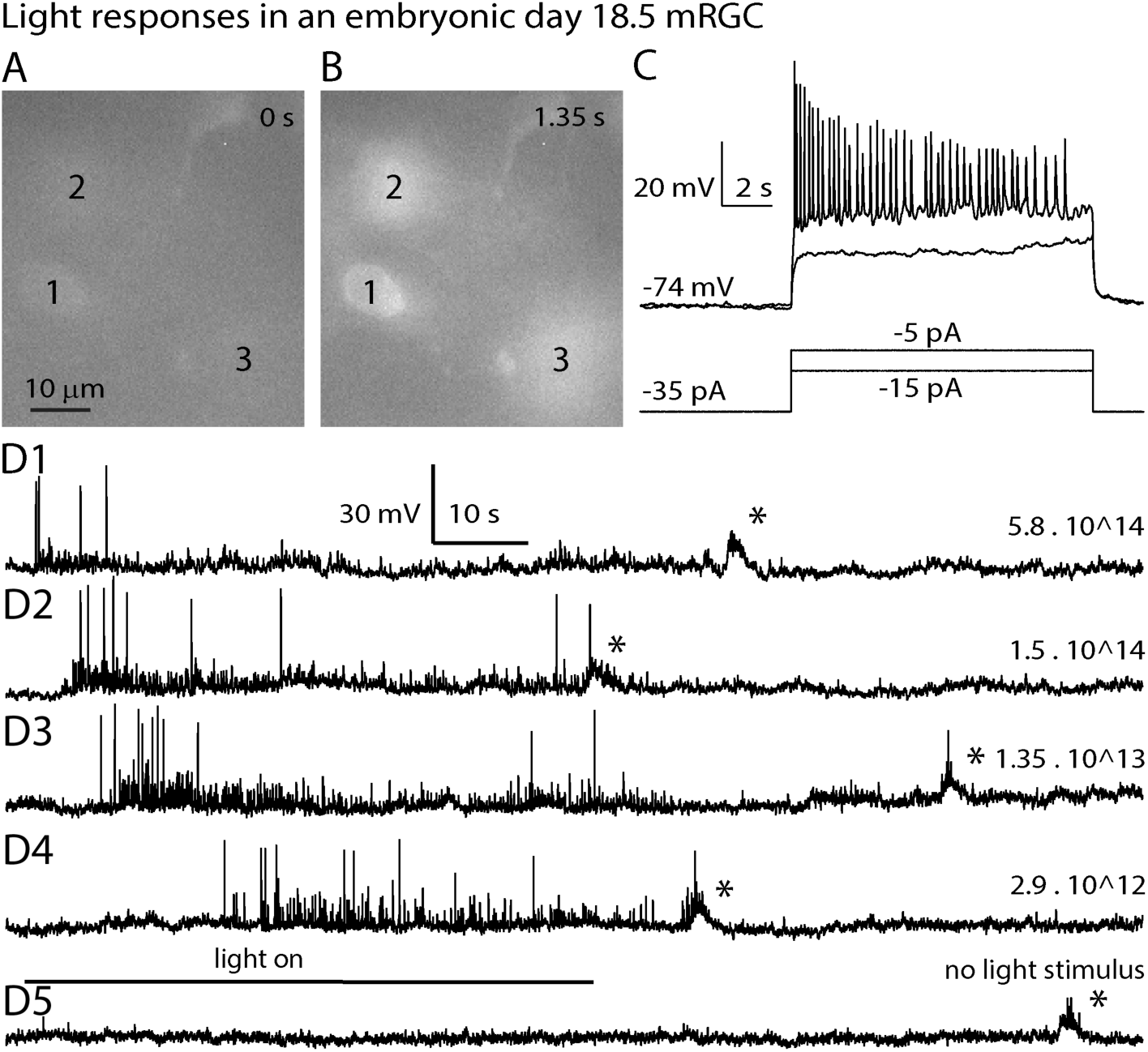
Light responses recorded from an mRGC in the retina of a foetal (E18.5) mouse pup. To select mRGCs for electrophysiological patch pipette recordings, we first surveyed the retina for light-induced calcium responses, which should be restricted to melanopsin cells (OPN4cre::Ai96). Then, using infrared Nomarski imaging, we searched for light-responsive mRGCs whose cell bodies were exposed on the inner surface of the retina. A and B show light-induced GCaMP6 increases in fluorescence in three cells. The images in A and B were obtained immediately after and 1.35 s after the onset of light stimulation, respectively. Although cells 1, 2 and 3 were light-responsive, the better-focussed cell 1 was targeted for electrophysiological recordings. C) Current clamp traces showing the responses of cell 1 to current injections to −15 and −5 pA from a holding current of −35 pA. The steps to −5 pA evoked repetitive spiking. D) Current-clamped light responses versus intensity from cell 1. The intensity of the 480-nm light stimulus, expressed as photons/cm^2^/s, is shown on the right. The bottom bar shows the timing of the light. The asterisks denote presumed spontaneous waves.

A population of low-amplitude spikes during light onset and occasional depolarizing “bumps” with a duration of approximately 1-2 s are evident in these recordings. We interpret the smaller spikes as spikelets conducted from neighbouring mRGCs as described by Arroyo et al. (2016). The bumps, which occur with a frequency of approximately 1/min, probably reflect the occurrence of spontaneous waves (Meister et al. (1990); Maffei L& Galli-Resta L (1990)).

### mRGCs in embryonic retinas are coupled via gap junctions

Cells in the adult mammalian retina are extensively coupled through gap junctions both to cells of the same type and to cells of different types. In the retinas of rats and mice, in particular, mRGCs are coupled to amacrine cells (Muller et al. 2010; Reifler et al. 2015). However, dye coupling between mRGCs in adult rodent retina has not been found, although Müller et al. (2010) and Ecker et al. (2010) specifically searched for homologous coupling between these cells. Precedent suggests that gap junction coupling could be age-dependent and that gap junction coupling may therefore exist in the retinas of younger mice. In spinal cord (Walton and Navarrete, 1991) and in the cerebral cortex (Kandler and Katz, 1991), gap junction coupling between neurons can exist early in development but is absent or much reduced in older animals. Consistent with a greater prominence of gap junction coupling in the retinas of very young animals compared to those of adults, Sekaran et al. (2003) reported that MFA, a putative blocker of gap junction coupling, reduced the number of light-responsive cells in early postnatal retinas. This was interpreted as evidence for gap junction coupling between mRGCs and other cells. Subsequently, Arroyo et al. (2016) found MFA-inhibitable functional and tracer dye coupling between genetically marked mRGCs in young neonatal mice (P4 to P8). We extended the work of Arroyo et al. (2016) to specifically test whether prenatal mRGCs are coupled to one another via gap junctions, pursuing the idea that younger mice may be more likely to express gap junction communication. We tested whether MFA, a compound that is commonly used to block gap junction coupling, reduced or eliminated light responses (Sekaran et al. 2003; Veruki et al. 2010). We examined the effects of MFA on light-evoked calcium elevations across a population of GCaMP-expressing mRGCs in fields of cells in flat-mounted retinas. Figure 10A shows the combined results obtained from 81 cells in two separate E16.5 retinas. MFA reversibly attenuated the amplitudes of the cells’ responses to light. The black dots in the top panel plot the response amplitudes after washout as a function of the response amplitudes observed prior to MFA application. The black dots are best fitted by a linear regression corresponding to y = 1.04 x. The data show good recovery from MFA. The blue dots plot the response amplitudes of each mRGC in MFA as a function of the response amplitudes observed prior to MFA application. The best-fitted linear regression line is y = 0.21 x. Non-parametric Wilcoxon Rank Sum Tests indicate that the reduction observed in MFA is highly significant (P = 8.3 × 10 ^−25^). These results are consistent with the spreading of light-evoked excitation between mRGCs via gap junctions. We performed the MFA experiments in 7 fields from 5 embryonic retinas. In 4 fields, MFA attenuated the responses reversibly. In the other 3 fields, the light responses recovered to more than 80% of their pre-drug levels in only a minority of the mRGCs. This apparent lack of reversibility is likely due to not having waited long enough after initiating washout (> 20 min). It should also be noted that in *OPN4cre::Ai96* mice we could only interrogate responses in mRGCs. Therefore, we were unable to use this method to rule out gap junction coupling to other non-melanopsin-expressing neurons. Additionally, a caveat associated with relying on MFA to identify gap junction coupling is that carbenoxolone, a previously used MFA-related compound, has been reported to attenuate L-type calcium currents in mRGCs (Bramley et al. 2011). Finding reduced numbers of light-responsive cells in MFA could also be attributed to the reduction of calcium currents.

**Figure 10.**
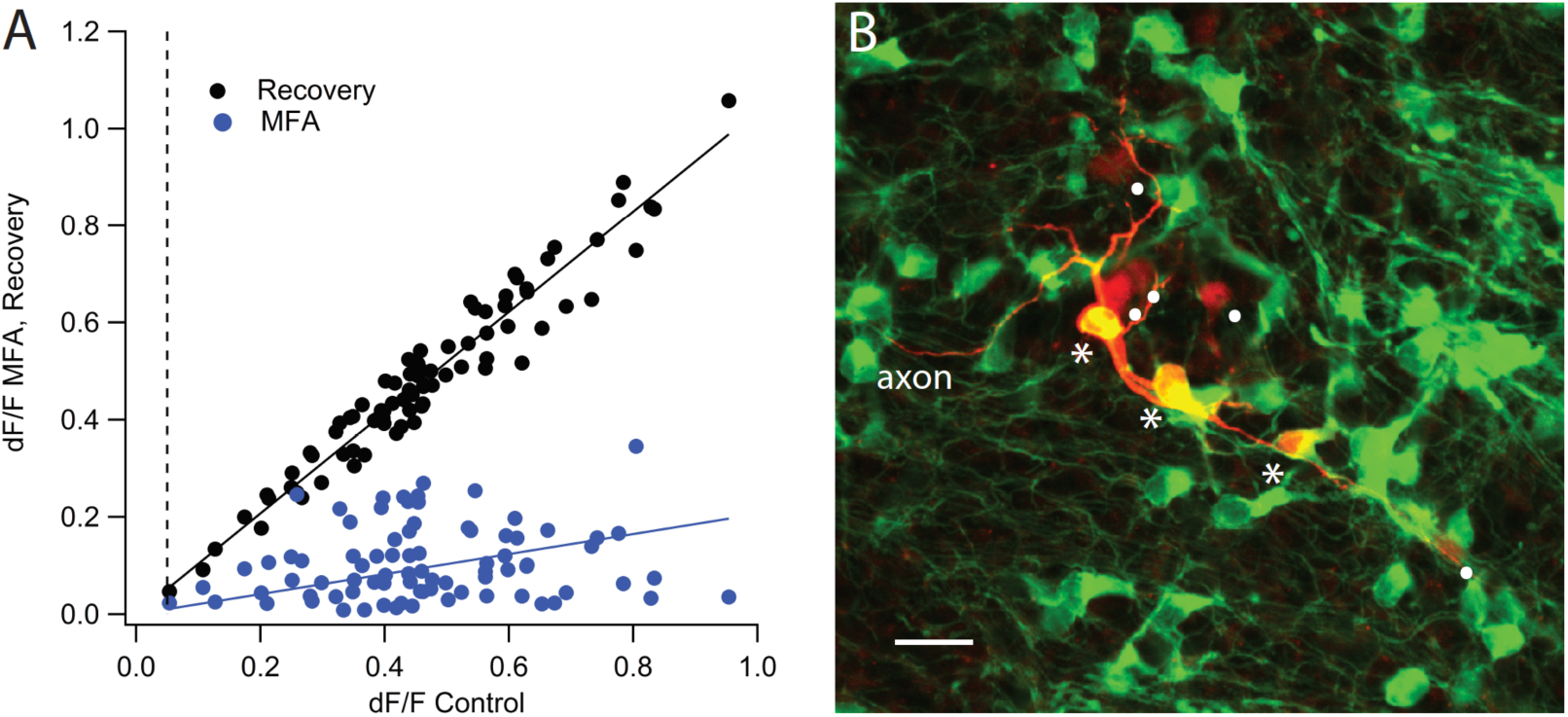
MFA, a gap junction-blocking drug, and intracellular neurobiotin injection reveal gap junction coupling between mRGCs in embryonic retinas. A) Meclofenamic acid (50 mM) strongly and reversibly reduces light responses in E16.5 mRGCs. (Top panel) Light-evoked dF/F of 81 fluorescent cells in 2 retinas containing mRGCs expressing GCaMP6 (OPN4cre::Ai96). dF/F in 50 mM MFA (blue dots) and following washout (black dots) was plotted versus control dF/F. The data include 81 of 142 light-responsive cells in which the response after recovery was at least 80% and no more than 120% of the control value (note that 142 cells responded to light onset under control conditions with dF/F’s > 0.05 but that in many of these cells the response failed to recover to at least 80% of the control value). The data were fitted with lines passing through the origin. The best fitting linear regression lines were y = 1.04 x and y= 0.21 x for recovery and MFA, respectively. The two-tailed P-value, obtained using the Wilcoxon Rank Test for paired data, was 8.3 × 10^−25^, indicating that MFA highly significantly attenuates light-evoked dF/Fs. MFA reversibly attenuated light responses in essentially all the mRGCs. B) Example of a neurobiotin injection (red) into an E18.5 mRGC. Three somas were neurobiotin- and EGFP-positive (green), indicative of tracer migration from one GCaMP6-expressing mRGC to at least two other mRGCs (stars). Five other somas (dots) were neurobiotin-positive (red) but lacked EGFP immunoreactivity, implying that mRGCs are coupled to non-melanopsin cells. The axon of the filled cell continued well beyond the picture to the left. The white scale bar indicates 20 μm.

To explore whether our MFA findings did indeed indicate coupling between mRGCs, we used patch pipettes to inject 17 embryonic mRGCs with neurobiotin as a means of tracing gap junction-mediated dye coupling to adjacent cells. Figure 10B shows an example of an mRGC that was injected with neurobiotin. Three somas were neurobiotin- and EGFP-positive, indicative of migration of the tracer from one GCaMP6-expressing mRGC to at least two other mRGCs (stars). At least five other somas were neurobiotin-positive (red) but lacked EGFP immunoreactivity. In summary, in 12 of 17 injections, at least one nearby mRGC cell was filled with neurobiotin. The maximum number of dye-coupled mRGCs observed to be connected an injected mRGC was 4. In 10 of the 17 injected mRGCs, the axons were filled with neurobiotin. Neurobiotin also filled from 0 to 7 non-mRGCs that were identified by the absence of OPN4cre-driven EGFP or tdTomato from their cell bodies. We were often able to follow neurobiotin-filled dendritic processes from the mRGC somas to the other dye-filled cells. These results provide strong evidence that mRGCs are coupled to other mRGCs as well as to other classes of inner retinal neurons in foetal mouse retinas.

## Discussion

The photopigment melanopsin is expressed in a subset of retinal ganglion cells termed mRGCs. Markers for melanopsin first appear in embryonic mouse and human retinas well before birth (embryonic day 11.5 and gestational week 8.6, respectively). mRGCs in postnatal retinas can be excited autonomously by light independently of visual inputs from rods and cones, the canonical photoreceptors in the eye. We recently demonstrated that exposure of pregnant females to light regulates the normal maturation of ocular vasculature in subsequently born pups. Based on our finding that the deletion of melanopsin in the foetal pups phenocopied darkness, we inferred that light stimulation of mRGC in the eyes of the foetuses modulates vascular maturation. There have been no reports directly demonstrating that mRGCs in foetal retinas are light-sensitive, raising some doubts about whether the ambient light in the mouse colony in which the pregnant females were housed was sufficient to excite mRGCs in the eyes of the foetuses. Here, using both intracellular calcium indicators and patch pipette recordings, we show that foetal mouse mRGCs respond to light as early as E15.5, four days before birth. We report further that foetal mRGCs share many of the phototransduction pathways that have been previously identified in mRGCs in the retinas of postnatal animals. Pharmacological experiments reveal that light responses are initiated by the coupling of light-activated melanopsin to a G_q/11_-G-protein cascade. We also show that calcium influx through L-type calcium channels and cation influx through TTX-sensitive sodium channels boost the amplitude of photoresponses. Electrophysiological recordings from individual embryonic mRGCs demonstrated that these cells can respond to light as dim as 2.9 × 10^12^ photons/cm^2^/s, providing strong supportive evidence that the ambient light in rooms where the pregnant females were housed is sufficient to excite mRGCs in foetuses. Additionally, we found that when the dye neurobiotin was injected into individual single mRGCs, it flowed into other nearby mRGCs via gap junction coupling. This finding contrasts with reports performed in adult rodent retinas in which no mRGC/mRGC coupling was discovered. Finding photosensitivity in the embryonic retinas of mice raises the question of whether human foetuses and early preterm infants could be sensitive to light and whether light exposure might modulate ocular development. Given that light stimulation of mRGCs plays a role in entraining circadian rhythms and mood in postnatal animals, some consideration should be given to the consequences of light exposure experienced by human foetal or preterm infants. Light exposure sources might include lighting regimens in neonatal care units and light exposure during intrauterine surgery. Such exposure might influence ocular development and/or behavioural states.

### No evidence was found that other photopigment-expressing cells in the foetal retinas of mice are light-sensitive

Could cells other than mRGCs in the embryonic eye confer light sensitivity that might affect ocular development? The present study focussed on the light sensitivity of mRGCs in retina. The results of the Calbryte 590™ experiments (Fig. 1) argue against the idea that OPN3- and OPN5-expressing cells generate light responses in the foetal mouse retina. Strictly speaking, we did not address the question of whether other non-retinal cell types that might express melanopsin prenatally could regulate ocular development. However, the preponderance of evidence presented here and in a previous study (Rao et al. 2013) strongly supports the idea that light stimulation of mRGCs is sufficient to account for light modulation of ocular vasculature maturation. In adult rodents, melanopsin expression has been found in the cornea, the iris and the retinal pigmented epithelium (RPE). We recently showed that melanopsin-containing fibres and cells in the corneas of postnatal animals were not light-responsive, suggesting that melanopsin might play a role in other sensory modalities such as thermal detection (Delwig et al. 2018). While light-sensitive melanopsin cells have been identified in the iris, there are no reports that these cells are light-sensitive in embryonic retinas or even that melanopsin markers are found in the embryonic irides. Melanopsin gene expression has been reported in cultured lines derived from RPE. There is no evidence that native RPE or cultured RPE cells lines are light-responsive. Further study focussed on the iris and RPE would be required to ascertain whether light stimulation of these cell types alters the vascular maturation of the eye.

### mRGCs are coupled electrotonically to one another in the foetal retina

Syncytial networks of electrotonically coupled neurons facilitate the spread of excitation across regions of neural tissue and coordinate the timing of excitability and rhythmic oscillations (Nagy et al., 2018). Electrotonic coupling between heterologous and homologous classes of neurons is common in both mammalian and non-mammalian neural tissues. Moreover, it is commonly found that electrotonic coupling is more prominent in the tissues of younger animals (Walton and Navarrete, 1991, Kandler and Katz, 1991). In the retinas of adult mice, electrotonic coupling of mRGCs to other mRGCs seems to be absent [Müller et al. (2010), Ecker et al. (2010)], but it is observed in younger postnatal mice (Sekaran et al. 2003; Arroyo et al. 2016). In the present study, we found electrotonic coupling between mRGCs. Additionally, we have evidence for electrotonic coupling from mRGCs to heretofore-unidentified non-mRGC cell types in the inner retinas of foetal mice. Electrotonic coupling between mRGCs and other heterologous neurons raises the possibility that light exposure during foetal and early postnatal maturation could excite neighbouring ganglion cells that project axons to the higher visual centres. In postnatal animals, light deprivation produced by suturing shut the eyelid of one eye disturbs the normal segregation of eye-specific targets in higher visual centres. This suggests that light excitation of mRGCs in neonatal animals before the normal age of eyelid opening may play a previously unrecognized role in retinotopic mapping in the higher visual centres.

Another notion drawn from the results presented here is that foetal light exposure might play a role in the formation of ocular dominance columns in primates. Horton and his colleague (Horton & Hocking, 1996) found ocular dominance columns in newborn macaques that were delivered by Caesarean section and kept in total darkness for a week after birth. Horton and Hocking argued that the formation of columns was genetically programmed based on the assumption that there had been no light-induced activity in the retinas of foetal monkeys. Given our findings in mice, it is worth considering the idea that ambient light might stimulate mRGCs in embryonic monkeys and thereby play a previously unrecognized role in the formation of ocular dominance columns.

### Light excitation of mRGCs in foetal retinas could modify patterns of retinogeniculate connections

Retinal ganglion cells extend axons to the lateral geniculate nucleus several days before birth. The postsynaptic contact regions in the LGN exhibit eye-specific contact regions. A consensus opinion is that spontaneous activity, often observed as propagated waves of activity, plays a critical role in segregation (Shatz CJ & Stryker MP (1988); Shatz CJ (1996); Meister et al. (1991); Maffei & Galli-Resta, (1990). Renna et al. (2011) and Arroyo et al. (2016) reported that light modulated the temporal structure of retinal waves via activation of mRGCs in the retinas of young neonatal mouse pups. Renna et al. (2011) also reported that mRGC excitation perturbed the eye-specific targeting of RGC axons in the LGN. Given our demonstration that light can activate mRGCs in foetal retina, we suggest that light exposure of pregnant females might play a role in establishing retinogeniculate connectivity in the maturing pups. Light could induce activity in mRGC axons targeting the LGN, or it could induce activity in neighbouring RGCs that might be electrotonically coupled to mRGCs. Another alternative is that mRGC excitation could modulate the structure of spontaneous waves that propagate across the foetal retina. Further studies will be needed to establish whether the light-induced activity of mRGCs as young as E15.5 is conducted to the LGN.

### Are human foetuses sensitive to light and might ambient lighting conditions at early gestational ages affect eye development?

Third-trimester human foetuses respond acutely to light directed through the mother’s abdominal wall. Examples of photic induced responses include modulation of heart rate and body movement (Kiuchi et al. 2000) and visually evoked brain activity (Eswaran et al. 2004, McCubbin et al. 2007). Since rods and cones begin functioning at approximately gestational week 35, the acute actions of light stimulation could be activating rods and cones as well as mRGCs. In any case, the occurrence of light-evoked responses in human foetuses demonstrates that light can excite photoreceptors in the foetal eye after passing through the abdominal wall of the pregnant mother.

Human eye development might also be sensitive to ambient light at gestational ages near the end of the first trimester. Human foetuses at gestational day 60 are at the same developmental age as mouse foetuses at gestational day 16.5 (Clancy et al. 2001), the age at which ambient light modulates subsequent vascular maturation in mice. Relevant to the possibility that reduced daily photoperiod might influence eye development in humans, an analysis of 276,911 conscripts of the Israeli Defense Forces between 2000 and 2004 found an association between an increased incidence of moderate and severe myopia and season of birth. Conscripts for whom the end of the first *in utero* trimester occurred during the winter months exhibited an increased risk of myopia (Mandel et al. 2008). One intriguing explanation is that light exposure of foetuses during the first trimester influence eye growth. While Mandel et al. (2008) ruled out family planning as a variable many other confounds need to be considered including diseases experienced during winter months, Vitamin D deficiencies, birth weights and other factors. In another analysis, Yang et al. (2013) reported that infants whose early gestation occurred during the winter months were at increased risk of severe retinopathy of prematurity (ROP). These findings raise the possibility that ambient light passing across the abdominal wall of pregnant mothers early in gestation might be considered as a factor that could affect eye development.

## Biographies

**Jan Verweij** acquired his doctorate in neuroscience at the University of Amsterdam under the mentorship of Martin Kammermans, where he studied synaptic and electrotonic interaction between rods and cones in goldfish retina]. He joined the laboratory of Denis Dacey at the University of Washington, where he studied synaptic interactions in monkey retina. Then, he joined the laboratory of Julie Schnapf at UCSF to study functional interactions between rods and cones in monkey retina.

**Shawnta Y. Chaney** received her B.S. in Biology and her Ph.D. in Biochemistry at the University of Houston, Houston TX. There, she studied the effects of gestational lead (Pb2+) exposure on mouse retinas. She further studied retinal development as a postdoctoral scholar in David Copenhagen’s lab at UCSF, where she focussed on embryonic and early postnatal light activation of melanopsin ganglion cells. Shawnta is now an Ocular Scientist at UNITY Biotechnology, where she tests therapies for age-related ocular diseases.

## Acknowledgements

We thank Anton Delwig and Michael Stryker for valuable discussion and advice during the course of this research. We thank Andrew Ishida for his critique of the manuscript. For technical advice, we thank Franklin Caval-Holme, James Bothelo III, Travis Porco and Jessica Wong.

